# Expectation-Based Gist Facilitation: Rapid Scene Understanding and the Role of Top-Down Information

**DOI:** 10.1101/2021.02.04.429791

**Authors:** Dominic McLean, Louis Renoult, George L. Malcolm

## Abstract

Scene meaning is processed rapidly, with ‘gist’ extracted even when presentation duration spans a few dozen milliseconds. This has led some to suggest a primacy of bottom-up information. However, gist research has typically relied on showing successions of unrelated scene images, contrary to our everyday experience in which the world unfolds around us in a predictable manner. Thus, we investigated whether top-down information – in the form of observers’ predictions of an upcoming scene – facilitates gist processing. Within each trial, participants (*N*=336) experienced a series of images, organised to represent an approach to a destination (e.g., walking down a sidewalk), followed by a final target scene either congruous or incongruous with the expected destination (e.g., a store interior or a bedroom). Over a series of behavioural experiments, we found that: appropriate expectations facilitated gist processing; inappropriate expectations interfered with gist processing; the effect of congruency was driven by provision of contextual information rather than the thematic coherence of approach images, and; expectation-based facilitation was most apparent when destination duration was most curtailed. We then investigated the neural correlates of predictability on scene processing using ERP (*N*=26). Congruency-related differences were found in a putative scene-selective ERP component, related to integrating visual properties (P2), and in later components related to contextual integration including semantic and syntactic coherence (N400 and P600, respectively). Taken together, these results suggest that in real-world situations, top-down predictions of an upcoming scene influence even the earliest stages of its processing, affecting both the integration of visual properties and meaning.

---

Apart from at waking, every environment we encounter is part of a progression of scenes unfolding around us as we move through our surroundings. As such, any single scene is not confronted in isolation but is instead simply the most recently perceived environment within the continuous experiential flow of our passage through the world. However, scene perception research has largely ignored such an asseveration, focusing more on the mechanisms responsible for processing segregated, individual scene images. In a traditional experiment, a participant may be faced with an image of a mountain, followed by a church, a kitchen, and so forth, a scenario clearly divergent from the progressive and structured environments one inhabits within the course of daily life.

We have, without question, learnt a great deal from investigation of the processing of isolated scene images, and such paradigms have been highly effective in identifying the mechanisms and visual features that facilitate processing of the initial meaning, or conceptual ‘gist’ (see Oliva, 2005), of a scene. Perhaps the most fundamental of findings is that this form of gist – the ability to derive the semantic information contained within a perceptual landscape – can be extracted even under conditions where viewing times span less than a tenth of a second (e.g., Potter, 1975). Such limited durations have led many to infer the primacy of bottom-up visual factors in rapid scene perception (Itti et al., 1998; Potter et al., 2014; Rumelhart, 1970), with the conviction that top-down information can have only a limited role under such brief time frames. This is a fair assessment if one contends that initial scene processing takes place in a classic hierarchical fashion. In such a scenario – of progressive activation through a linear pathway of anatomical areas divergent in terms of functional specificity – it is unlikely top-down feedback would be received prior to such rapid scene categorisation taking place. Therefore, while such models do not deny the role of feedback or re-entrant connectivity as processing continues through time, they propose feature-extraction mechanisms as sufficient for distinguishing conceptual information and meaning within complex natural scenes, with a single ‘forward sweep’ of neural activity through the ventral stream (Potter et al., 2014).

However, the traditional view of the serial processing of visual input has been questioned for some time (Engel et al., 2001; Ullman, 1995), and the latest recurrent models can better explain human visual recognition when compared to feedforward neural networks (e.g., Spoerer et al., 2020). Likewise, the past decade has seen great advances in our understanding of the broad extent of reciprocal connections within the neural architecture (e.g., Groen et al., 2017; Kravitz et al., 2013). For instance, research methodologies spanning MEG, EEG and TMS have all provided evidence for rapid local recurrent processes within early visual cortex (Boehler et al., 2008; Camprodon et al., 2010; de Graaf et al., 2014; Foxe & Simpson, 2002), with the proposal that these processes might start only a few tens of milliseconds after the arrival of the visual input (de Graaf et al., 2012). Furthermore, multiple feedforward-feedback loops have been hypothesised as taking place within the first 100 ms of stimulus onset (Bullier, 2001; Juan & Walsh, 2003), a proposition strengthened by the finding of activation in intermediate visual areas prior to the completed contribution of early visual cortex (Koivisto et al., 2011).

Similarly, from the object processing literature, evidence reveals that top-down processes are initiated prior to completion of target recognition, with the suggestion that early activation of higher-order brain regions facilitates the systematic analysis of bottom-up information (Bar et al., 2006). In other words, low spatial frequency information is passed rapidly to higher areas and is then used to form predictions as to the identity of the object being viewed. Consequently, this allows for the pre-activation of a limited set of object representations which are subsequently matched against the continuing flow of bottom-up information (Bar et al., 2006). It seems reasonable to infer that some equivalence may exist within the manner of operation for scene processing, whereby an initial ‘sketch’ (Marr, 1982; Rensink, 2000) of the environment may allow for the pre-activation of scene representations in higher-order areas. Indeed, parallel co-activity within higher regions has been observed even while perceptual coding of visual scenes is actively proceeding (Catherwood et al., 2014). While concurrent activation cannot be taken as direct evidence for interaction between regions, it provides the opportunity for such interactions to a far greater extent than models which assume somewhat step-by-step activation, whereby higher-order processing occurs only as perception subsides.

The above research shows that the selection and processing of those elements within even the precursory stages of the feed-forward wave of activity may be open to facilitation. Moreover, if top-down information can rapidly influence bottom-up processing in scenarios such as these – where no indication as to what will be displayed is provided prior to stimulus onset – then it seems appropriate to suggest that top-down influence might be even more rapid when pre-target cues allow for a subsequent visual image to be predicted. Such a claim is reinforced when considering the growing weight of evidence demonstrating that activity within the visual cortex, including early striate cortex, can be affected by expectations alone (e.g., Aitken et al., 2020; Grill-Spector & Malach, 2004; Kok et al., 2012), that the shape-selectivity of neurons in area V1 is altered depending on what geometric shape is expected (McManus et al., 2011), and that *a priori* expectations generated by scenic context can lead to increased activation in higher-order areas during subsequent visual processing (Caplette et al., 2020). Accordingly, here we present an investigation as to whether an observer’s expectations of an upcoming scene category have a direct effect on the initial stages of processing, i.e. the extraction of conceptual gist. In so doing, we attempt to better replicate how scenes are processed outside the laboratory, namely as predictable settings preceded by contextually relevant visual information, and hence proffer that models based on a progression of activation across successive regions cannot provide an exhaustive account of functionality.

In concordance, considerable evidence signals that expectations can influence subsequent processing of the environment, such as the inadvertent bypassing of crucial but unexpected visual information (Mahon, 1981), ‘looked-but-failed-to-see’ traffic accidents (Langham et al., 2002), and increased task-related errors in situations inconsistent with expectations (e.g., Endsley & Garland, 2000). Relatedly, the influence of context-based expectations on cognitive processing has been widely investigated through the experimental manipulation of object-scene relationships. Such research has repeatedly shown that target objects are found more quickly (Biederman et al., 1973; Võ & Henderson, 2011), and with higher accuracy (Antes et al., 1981; Davenport & Potter, 2004; Underwood, 2005), when within ‘appropriate’ scenes (i.e., where the scene category and target object are semantically congruous). In addition, such context effects have been found not only during the simultaneous presentation of a scene and target object, but also when a scene image is presented prior to (Demiral et al., 2012; Ganis & Kutas, 2003; Võ & Wolfe, 2013), and independent of (Palmer, 1975), object presentation. Due to the speed with which objects can be detected and identified (e.g., Crouzet & Serre, 2011; Kirchner & Thorpe, 2006; Thorpe et al., 1996), these studies demonstrate that semantic information can rapidly influence visual processing, and also that increased processing ability related to congruency is evident even when natural scene images are used as a precursory means of inducing expectations. If scenes can provide semantic information capable of altering subsequent object processing, it would seem intuitive that such influence similarly extends to subsequent scene processing.

Indeed, experimental evidence has demonstrated that a scene can be primed by a preceding scene-image, termed the ‘scene priming’ effect, although this has largely concerned priming at the perceptual – rather than conceptual – level. Increased performance regarding spatial layout judgements have been elicited when target scenes are primed using an identical scene image (Sanocki, 2013) or with images of the target scene from different viewpoints (Sanocki & Epstein, 1997, although see Epstein et al., 2005), while image detection ability is improved if primed across scenes more closely matched in terms of spectral information (Caddigan et al., 2017). Furthermore, when primes and targets are adjacent segments of the same complete landscape – thus intrinsically different while being similar in general composition – biases to cortical responses, alongside improved feature detection performance, have been shown (Blondin & Lepage, 2005). However, the mechanisms behind such effects are open to debate, as much of this work is proposed to reflect the maintenance of scene layout information in memory (Oliva & Torralba, 2001) or simply the priming of low-level visual features (Brady et al., 2017; Shafer-Skelton & Brady, 2019; although see Sanocki, 2013 for a potential top-down explanation). The focus of the current study, on the other hand, is investigation of the effect of expectations on scene processing at the semantic level. It is, therefore, equivalent to conceptual (Tulving & Schacter, 1990) or semantic priming (Meyer & Schvaneveldt, 1971), and so more similar to research showing performance increases when a scene’s category membership is presented in text prior to presentation of the target image (Reinitz et al., 1989).

Correspondingly, recent research has further suggested a potential influence of top-down factors over the limited duration of gist processing (Greene et al., 2015). Here, briefly presented atypical scenes – such as a boulder in the centre of a living room, or a pillow-fight in a town square – were found to be more difficult to both process and understand compared to frequently encountered scene types (e.g., a car in a driveway). This indicates that an observer’s prior semantic knowledge can influence the rapid processing of complex natural scenes, even over highly curtailed presentation durations. However, the design of that study still involved the presentation of single, unrelated images on each trial, and so cannot apprise us of the interaction between immediately preceding information and predictability. So, while such research highlights the cost of violating the expectations held in long-term memory, it speaks less to the violation of expectations built upon the ‘on-line’ flow of information as it is received.

This gap in understanding needs addressing due to how we experience the world around us, where the daily sequential emergence of scenes takes place in a predictable fashion. This predictability is not only apparent for locations with which we are familiar, such as knowing what scene will greet us when turning the corner of a street travelled daily, but is also related to our expectations when in previously unencountered locations. When walking down an unfamiliar street, in an unfamiliar town, experience with similar environmental surroundings allows one to form predictions as to what awaits past the next corner. The sight of houses at the end of the street may fit within the expected sequential flow of situational contexts built over a lifetime of similar experiences, thereby allowing for efficient cognitive processing (Bartlett, 1995). The sight of a volcano, on the other hand, would most likely violate any such schema (e.g., Mandler & Ritchey, 1977), resulting in the allocation of greater cognitive resources in order to process such unexpected information (Barlow, 1961; Haque et al., 2020).

Recent research has started to address this directly, by pointing towards the influence of predictions on gist processing through the use of pre-target narrative sequences (Smith & Loschky, 2019). Here, the spatiotemporal coherence of image sequences depicting different routes (such as from an office to a parking lot) was manipulated. When image sequences were presented in narrative order, as opposed to when randomised, categorisation performance for – and predictability of – target scenes was significantly increased. While this work was concerned with the ordering of pre-target images, rather than their congruency with an upcoming target-scene, it reveals that expectations as to what scene may be encountered next can be informed by what has gone before and, moreover, that these expectations may have a functional role in terms of facilitating scene-gist processing. An explanation for the underlying mechanisms has been offered, whereby narrative sequences help construct a current event model, which then in turn influences the extraction of gist information (Smith & Loschky, 2019). As a consequence, an iterative process is created whereby ‘front end’ information extraction (such as that derived from attentional selection mechanisms) informs ‘back end’ model construction (initially stored in working memory), which in turn influences front end processes, and so forth (Loschky et al., 2019).

So, both directly and indirectly, previous work has indicated that observer expectations can affect scene gist processing. Of equal importance, such a suggestion does not seem unreasonable when considering the typical mechanisms of visual processing more broadly. While we exist within a world of seemingly limitless sources of sensory information the visual system is constrained by limited processing capacity, and so it has long been understood that increased efficiency can be derived through drawing on learned experience to aid our interaction with the environment (Chaumon et al., 2008; Fiser et al., 2016; Gregory, 1997; Li et al., 2004; Roc, 1997; Ullman, 1980; although see Gibson, 2014 for an account of ‘direct’ perception). With this in mind, for the visual system not to use expectations to facilitate scene gist processing would seem to contravene its typical mode of operation.

The emergence of predictive coding models provides a potential framework by which the generation of expectations as to upcoming visual stimulation might, in part, offset inherent signal transmission delays (Hogendoorn & Burkitt, 2018; Nijhawan & Wu, 2009; Rauss et al., 2011). While it is beyond the scope of the current study to make determinations as to the precise mechanisms involved in any top-down influence on gist processing, such models provide a viable solution. For example, any current perceptual environment may lead to predictions of the subsequent environment, resulting in the pre-activation of those internal representations. These expectations may subsequently influence early visual areas by adapting their processing of perceptual features, through adjustment of prediction error thresholds, based on the representations chosen as likely to fit the upcoming landscape (Rauss et al., 2011). As such, the neural signal pattern even at early stages of the processing stream might be a reflection of a perceptual landscape’s congruence with predictions, above-and-beyond merely a reflection of the low-level information contained within (Mumford, 1992).

To tease apart the role of on-line expectations within processing, the current study investigated the influence of visual information received immediately prior to target-scene onset. Across all experiments we employed a fundamental change to the traditional methodologies, which either position targets within a rapid serial visual presentation (RSVP) sequence of unrelated images (e.g., Potter, 1975) or present only a single image per trial (e.g., Greene et al., 2015). This was achieved by providing contextual information through presentation of antecedent ‘lead-up’ images, allowing us to investigate the influence of just-prior experience on the understanding of scenes. These leading images provided a flow of movement through an environment and towards a scene, and so represented an approach to a destination. This is, we suggest, a more naturalistic means by which to generate predictions based on lifelong experience, and as a result is somewhat removed from research investigating the effect of predictions on perception using simplistic pre-target cues (e.g., Summerfield & Koechlin, 2008), or where predictability is manipulated by synthetic means such as the learning of arbitrary contingencies prior to task commencement (e.g., Hindy et al., 2016). A key aim of the current study, therefore, was to provide a more ecologically valid reflection of scene perception. While only an approximation of this can be achieved with a sedentary participant viewing static images on a monitor, careful construction of image-series was considered sufficient in affording an impression of progress through a landscape.

Then, by manipulating whether the target scene was congruous with these leading images, i.e. the ‘approach-destination’ congruency, we hoped to demonstrate whether there is indeed an influence of predictability on scene categorisation ability. In addition, across the separate behavioural experiments we manipulated the presentation duration of destination images, the spatiotemporal coherence of approach-image sequences, and the provision of pre-destination scenic context in order to more fully investigate the mechanisms underlying the effect of expectations on gist processing. Finally, we turned to electroencephalography to map changes in brain activity relating to the manipulation of approach-destination congruency, with the aim of identifying the forms of cognitive processing most readily affected by the violation of expectations.

## Experiment 1a

The ability to categorise scenes even under the briefest presentation durations has led many to argue that such rapidity of processing must take place largely outside the involvement of top-down influence (Itti et al., 1998; Potter et al., 2014; Rumelhart, 1970). On the other hand, more recent research has found that semantic information can influence scene processing within shorter timeframes than previously thought (Greene et al., 2015; Võ & Wolfe, 2013). However, this research has largely focused on the semantic congruity of objects within a scene, rather than congruity between scenes. We aimed to address this gap by presenting series of ‘approach’ images prior to ‘destination’ target scenes, while manipulating the congruency between the destination and its forerunners, in order to investigate whether semantic predictability of an upcoming scene influences processing.

Furthermore, in Experiment 1a we manipulated target presentation duration to investigate whether the influence of contextual information remained consistent across the different stages of scene processing. Specifically, models assuming primacy of bottom-up factors during gist processing would not expect differences in categorisation performance as a function of congruency at target durations below 100 ms. Under such models (e.g., Potter et al., 2014), the category of the lead-up scenes would be expected to have minimal influence during the gist processing of the subsequent target image. Conversely, if performance differences were found at such brief durations this would lend support to the proposition for top-down influences on gist processing.

We hypothesised that destination scenes preceded by congruous approach images would be more accurately categorised, compared to those with incongruous approaches. Additionally, we predicted that this benefit would be most apparent at briefer presentation durations. This was due to our expectation that, at shorter durations, the ability to extract visual information would be most curtailed whereas, at longer durations, enough visual information would be extracted and processed from destination scenes as to bring categorisation accuracy for all targets towards ceiling performance. Hence, we expected to see the biggest congruency-related differential in performance at the briefest target durations, as this would be the point of maximal benefit from providing participants with a congruous scenic context prior to destination onset.

### Design

All experiments were programmed and presented using PsychoPy (www.psychopy.org) version 1.85.3, unless otherwise stated (Peirce & MacAskill, 2018). An experimental trial began with participants viewing a sequence of five leading images, organised to represent an approach to a location. These approach images were followed by a target scene, representing a destination, which required a categorisation judgement from six available choices. All series depicted travel on foot, in order to convey a sense of walking through an environment (see Appendix A for additional details relating to the construction of series). Each participant sat 120 trials, 75% of which had leading images congruous with the target scene. This ratio was chosen to ensure participants remained attentive to the leading images. Target scenes could be from one of 30 separate categories, split equally between interior and exterior sceneries (see Appendix B for a list of categories used). Indoor and outdoor scenes vary from one another on fundamental characteristics such as level of expansiveness and roughness of textures (e.g., Oliva & Torralba, 2001), and there are suggestions that categorisation performance might differ across these two superordinate categories (Fei-Fei et al., 2007). Therefore, we chose to include both types of environment to provide a more complete picture of gist processing within typically encountered locations. All categories were considered familiar (e.g., ‘bathroom’, ‘beach’, etc.). Further to this, we manipulated target duration as a between-subjects variable, in order to investigate potential changes over the time-course of gist processing. Targets could be presented for 33, 50, 100 or 250 ms (2, 3, 6 or 15 frames on a 60Hz monitor). See Figure 1 for a schematic of the experimental protocol.

**Figure 1.**
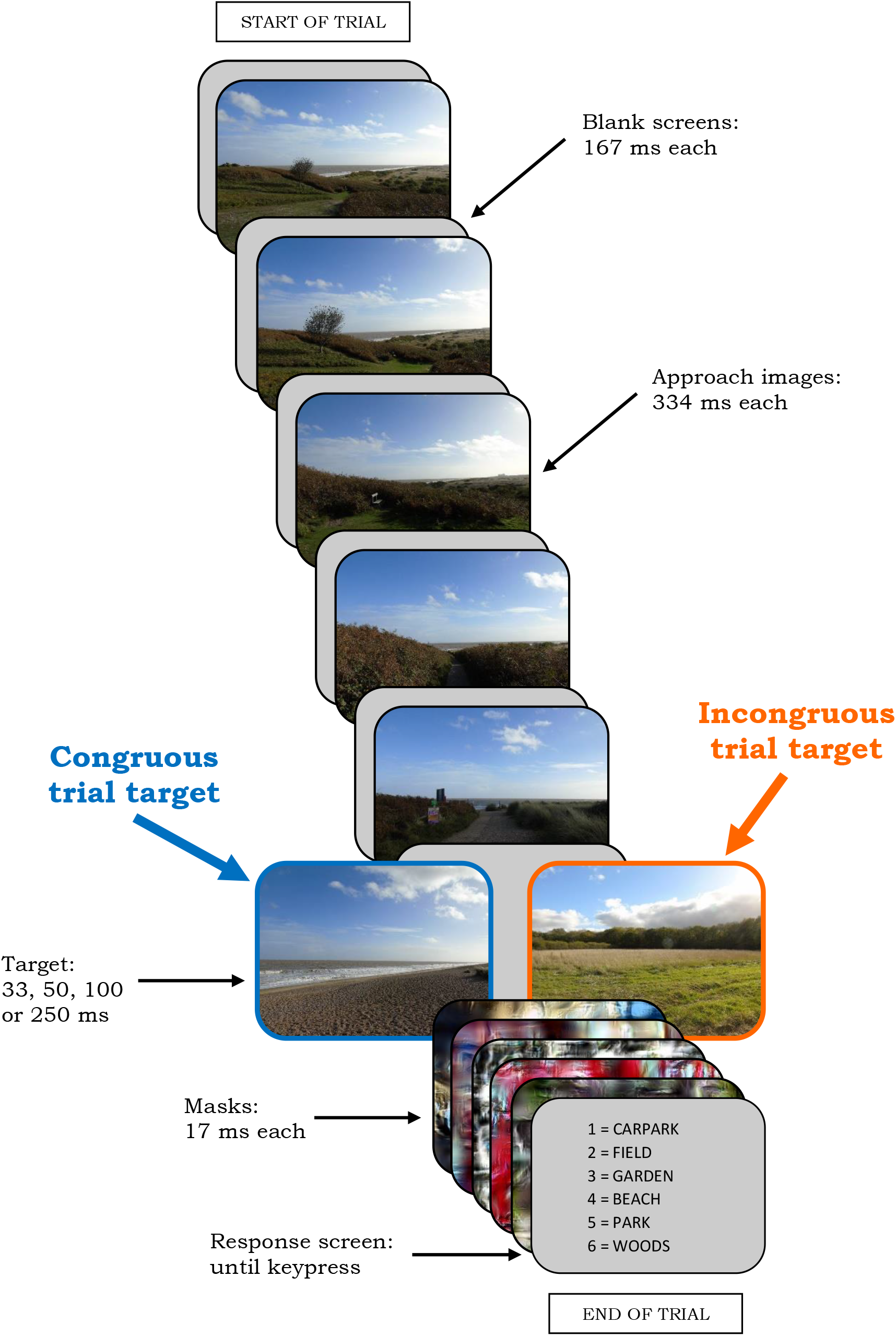
Schematic of the Protocol for Experiments 1a and 1b

### Participants

Our initial study included 129 undergraduate psychology students, recruited through the University of East Anglia’s research pool, who received course credits for participating (*M*_age_ = 20.09, *SD*_age_ = 3.68; 103 Females, 26 Males; 113 Right-handed, 15 Left-handed, 1 Ambidextrous). All experiments were approved by the ethics committee at the University of East Anglia’s School of Psychology, and all participants provided written informed consent prior to taking part in the study.

### Stimuli

The collection of images was comprised of photographs taken by the researchers alongside high definition images of sceneries and video-stills freely available on the internet. A total of 756 images were used as stimuli, of which 720 appeared in the experimental trials. No images were repeated. Each trial consisted of five spatiotemporally coherent approach images, followed by a target scene, resulting in 120 individual series. There were four series for each of the 30 scene categories. Approach images were sequential, first-person viewpoints heading towards a specific destination, with the aim of imbuing in participants a sense of progression through an environment.

One series from each of the scene categories was selected at random to become an Incongruous trial. The target scenes of each of these 30 series were then randomly reallocated amongst each other. This redistribution was conducted in adherence to two principles. Firstly, a target could not replace another target of the same scene category as, although it would be a different exemplar than what might be expected, it would still be semantically related to the approach images. Secondly, a target could only replace another target of the same superordinate category (in terms of interior / exterior distinction). This division was maintained due to suggestions that discriminating between superordinate categories is not analogous to discriminating between basic-level categories. While there is still debate as to the exact order with which these different levels are processed (see, for example, Banno & Saiki, 2015; Fei-Fei et al., 2007; Kadar & Ben-Shahar, 2012; Loschky & Larson, 2010), it was considered necessary to follow this principle to avoid potential changes in processing strategy from trial to trial. Each target image was followed by a set of five masks, presented rapidly in sequence. A different set of masks was used after each target. To achieve this, 600 masks were generated from the approach images by using Portilla and Simoncelli’s (2000) texture synthesis algorithm in Matlab, in line with previous research showing this to be an effective method for placing temporal constraints on bottom-up processing of scene images (Evans et al., 2011; Greene et al., 2015).

Performance was judged through participants selecting the category that best described each destination scene from a list of six options. The available category options on each response screen were allocated randomly and were also randomised in terms of item position. All options were of the same superordinate category (indoor / outdoor) as the target. This was to ensure participants could not reject certain options based simply on superordinate-level membership. For Incongruous trials, the category the approach images would be expected to lead to was also included. In other words, an Incongruous trial displaying an approach to a ‘Park’ followed by a ‘High Street’ destination would have a response screen that included ‘Park’, ‘High Street’ and four other exterior scene categories. All images and masks were displayed with an image resolution of 800 x 600. All images were presented in colour, on a monitor with a refresh rate of 60Hz.

### Procedure

Prior to starting the experiment, a series of instruction screens were displayed explaining the task and prompting participants to imagine travelling through the environments that were presented. This was followed by six practice trials, with the opportunity to ask any questions of the researcher on their completion. The same set of practice trials, in the same order, was experienced by each participant.

The 120 trials were presented in a different randomised order for each participant. Each trial included five sequential approach images, separated by blank screens, followed by a destination image. A series of five masks began at target-offset, prior to a 6AFC response screen. Once a response had been given, by pressing the number on the keyboard corresponding to the chosen category, the next trial began. At three equally spaced points within the task an ‘optional break’ screen was displayed, where participants could choose to pause if they wished and recommence once any key was pressed.

## Results and Discussion

One participant was removed from the analysis due to a zero score for the Incongruous condition, suggestive of a misunderstanding of the task. A further five participants were removed due to a score in either congruency condition being outside 3 standard deviations of the mean for the respective target duration. Analysis was conducted on the remaining 123 participants (*M*_age_ = 20.11, *SD*_age_ = 3.76; 97 Females, 26 Males; 107 Right-handed, 15 Left-handed, 1 Ambidextrous). The proportion of correct answers on both congruency conditions was calculated for each participant, and the mean scores across participants were plotted (see Figure 2). As can be seen, when the opportunity to extract visual information was limited due to brief target durations (33 and 50 ms conditions), performance on Incongruous trials was discernibly below that on Congruous trials. As the target duration increased, however, this disparity across congruency conditions narrowed (100 ms), and subsequently disappeared (250 ms).

**Figure 2.**
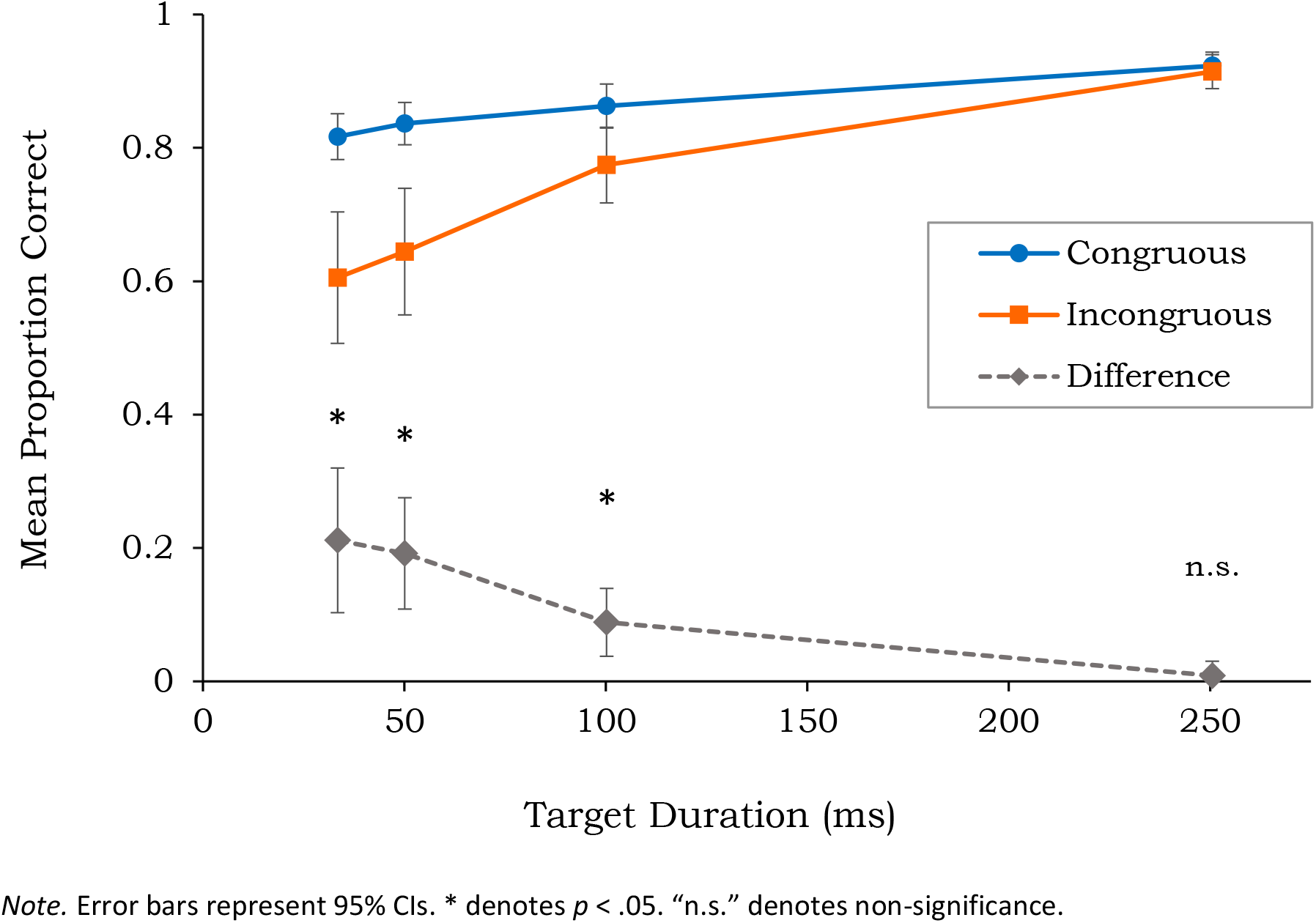
Scene Categorisation Accuracy for Congruous and Incongruous Trials as a Function of Target Duration

For each participant, accuracy scores for Incongruous trials were subtracted from scores for Congruous trials, to assess performance differences across congruency conditions. The values were first used to test the data for normality, which was found to be positively skewed at the 33, 50 and 100 ms target durations, and also to be leptokurtic at 100 ms. The non-normality for these three conditions was confirmed through Shapiro-Wilk tests (all *p*s < .005), and for this reason non-parametric alternatives were chosen for the analysis.

A Kruskal-Wallis test showed the differences in congruency-related scene categorisation accuracy to be significantly affected by the presentation duration of target sceneries, *H*(3) = 17.46, *p* = .001. Further to this, pairwise comparisons with Bonferroni-adjusted p values showed the disparity in performance relating to congruency differed significantly across target durations. The disparity in performance at a target duration of 33 ms was significantly different to that at 250 ms (*p* = .004, *r* = 0.44), and likewise at 50 ms compared to 250 ms (*p* = .001, *r* = 0.50). The difference between the 100 ms and 250 ms conditions only approached significance (*p* = .087, *r* = 0.32), and no other significant differences were found across durations.

Follow-up Wilcoxon tests were then employed to investigate potential congruency-related differences in categorisation accuracy at each target duration. Difference scores (Congruous accuracy minus Incongruous accuracy) were compared to zero at each duration using a one-sample Wilcoxon test. Bonferroni adjusted p values are reported. At 33 ms, participants were significantly more accurate at categorising Congruous trials (*Mdn* = 0.83) than Incongruous trials (*Mdn* = 0.73), *Z* = −3.74, *p* < .001, *r* = −0.47, representing a medium-to-large effect. The same pattern was true at 50 ms, with greater accuracy for Congruous trials (*Mdn* = 0.86) than Incongruous trials (*Mdn* = 0.77), *Z* = −4.39, *p* < .001, *r* = −0.54, representing a large effect. This was again apparent at 100 ms, with greater accuracy for Congruous trials (*Mdn* = 0.89) than Incongruous trials (*Mdn* =0.80), *Z* = −3.40, *p* = .003, *r* = −0.43, representing a medium effect. However, no congruency-related differences were found at 250 ms (*p* = 1), with similar accuracy scores for both Congruous (*Mdn* = 0.94) and Incongruous trials (*Mdn* = 0.93).

As predicted, in Experiment 1a we found a significant benefit to categorisation performance when a target scene was preceded by semantically congruous approach images, revealing that participants’ expectations were influencing scene processing. Furthermore, the greatest differential in performance across congruency conditions was seen at the briefest target durations (33 and 50 ms), indicative of gist extraction being modulated by top-down information. These findings sit in agreement with previous reports of expectations influencing subsequent processing of the environment (e.g., Langham et al., 2002), as well as research showing that object processing can be facilitated if situated within semantically compatible sceneries (e.g., Underwood, 2005). While facilitation of processing across scene-images has previously been observed in relation to the priming of visual features (Brady et al., 2017), we suggest that the benefit of congruency seen in Experiment 1a was due to the provision of semantically relevant context, and so more akin to the semantic priming of a scene when preceded by a relevant written word (Reinitz et al., 1989), or to work finding a disruption to gist processing when an observer views improbable sceneries (Greene et al., 2015). Therefore, these results revealed that top-down information – in the form of expectations generated prior to target scene appearance – was able to influence gist processing, a proposition at odds with models assuming minimal higher-order modulation of gist processing (e.g., Itti et al., 1998; Rumelhart, 1970).

## Experiment 1b

The results from Experiment 1a demonstrated an advantage in categorisation performance for Congruous trials, apparent at target durations where the opportunity to process visual information was most limited. However, it was important to confirm that these findings were due to the congruency manipulation as opposed to unintended residual effects based on the experimental design. Specifically, 75% of trials were congruous in Experiment 1a, and so higher performance on these trials was feasibly based on their increased frequency compared to Incongruous trials. We addressed this possibility in Experiment 1b, by switching the relative presentation frequencies of the congruency conditions.

The reduction in the number of Congruous trials in Experiment 1b also served a further purpose: in a task where most trials are incongruous it is not beneficial for participants to take account of the contexts provided by the approach images, as these are more often than not unrelated to the destination. If a pattern of results similar to those from Experiment 1a emerged, therefore, in terms of higher performance for Congruous compared to Incongruous trials, this would suggest that predictions as to an upcoming scene category were being generated automatically.

### Design

We again employed a 3:1 split across trial congruency, but now with 75% of trials having destinations incongruous to the approach images. The decision was also taken to limit Experiment 1b to three target-duration conditions. This was due to the preceding iteration showing very similar levels of performance, in terms of both congruency conditions, across the 33 and 50 ms target durations. There was also a noticeable amount of variation in performance across participants at 33 ms, with some failing to achieve scores above chance level. As a result, it was decided that the 50 ms condition provided the most reliable reflection of general performance under circumstances of limited availability of visual stimulation.

Although the selection of Incongruous trials in Experiment 1a had been achieved by random assignment, it was prudent to ensure this had not led to any bias through unintentional systematic differences across the two congruency conditions. As such, Experiment 1b introduced a Latin Square design. Four separate versions of the protocol were programmed, each with a different set of 30 Congruous trials (one from each scene category). This meant that, over the course of the experiment as a whole, all series were presented in both congruous and incongruous fashion, with the specific makeup of conditions determined by which version a participant sat. Versions were cycled through for each new participant, separated by target-duration condition.

### Participants

Our second experiment included 90 undergraduate psychology students, recruited through the University of East Anglia’s research pool, who received course credits for participating (*M*_age_ = 20.89, *SD*_age_ = 4.98; 68 Females, 22 Males; 81 Right-handed, 9 Left-handed).

### Stimuli

Experiment 1b used the same image set and masks as Experiment 1a. The response screens were redrawn, using the same randomisation procedures as the first experiment.

### Procedure

The procedure for Experiment 1b mirrored that of 1a, with one alteration. A handful of participants had asked for clarification of certain category words during the previous iteration, most notably ‘Quay’. To eliminate this issue, prior to beginning Experiment 1b participants were shown a list of the 30 scene categories and were provided with explanations by the researcher where needed. Participants were assured that the list did not need to be memorised.

## Results and Discussion

Four participants were removed due to a score in either congruency condition being outside 3 standard deviations of the mean for the respective target duration. Analysis was conducted on the remaining 86 participants (*M*_age_ = 20.98, *SD*_age_ = 5.08; 65 Females, 21 Males; 77 Right-handed, 9 Left-handed). The proportion of correct answers on both congruency conditions was calculated for each participant, and the mean scores across participants were plotted (see Figure 3). As in Experiment 1a, when the opportunity to extract visual information was limited due to a brief target duration (50 ms), performance on Incongruous trials was some distance below that on Congruous trials. As target duration increased, the disparity across congruency conditions narrowed (100 and 250 ms).

**Figure 3.**
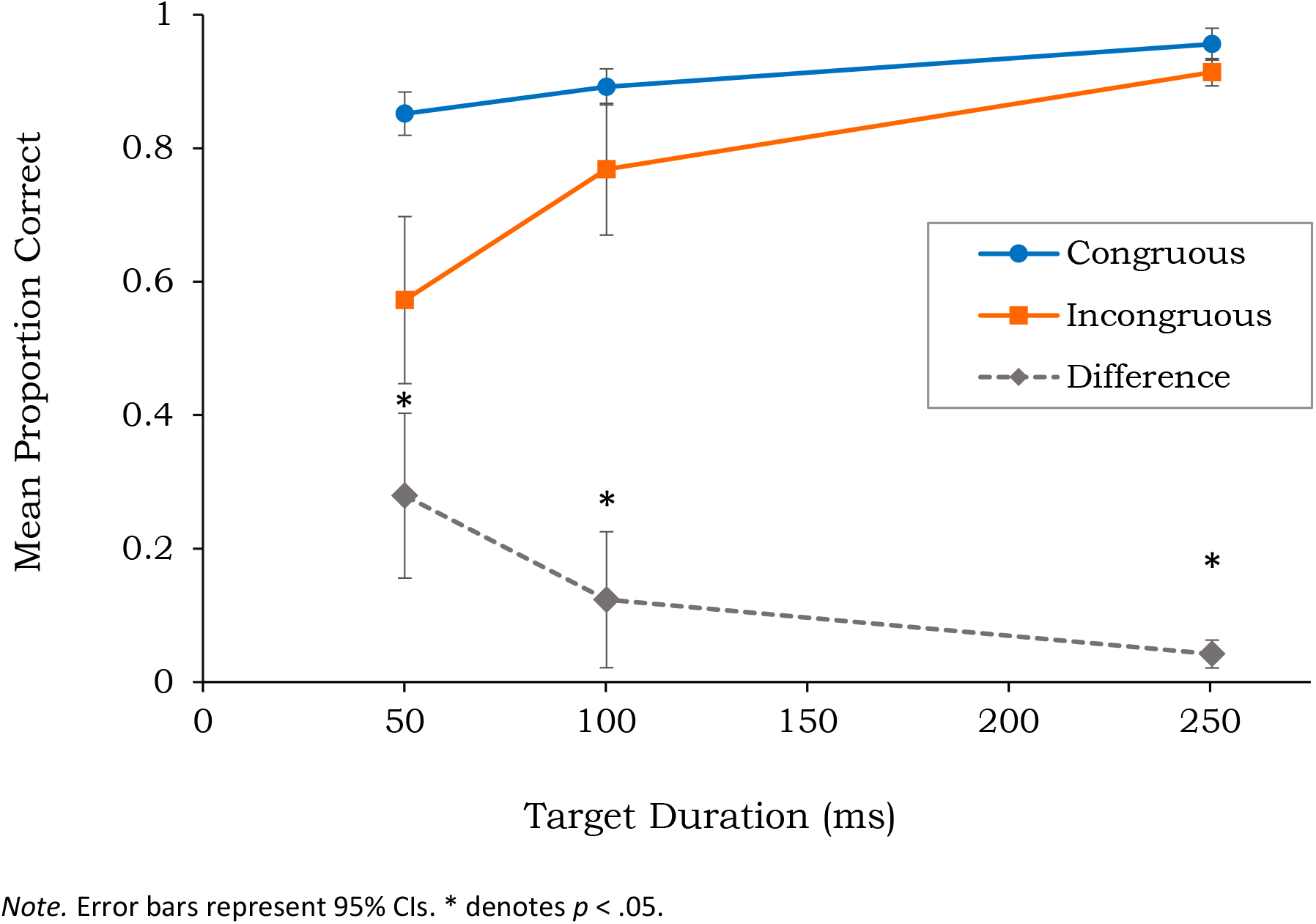
Scene Categorisation Accuracy for Congruous and Incongruous Trials as a Function of Target Duration

Accuracy scores for Incongruous trials were subtracted from scores for Congruous trials, to assess performance differences across congruency conditions. The values were first used to test the data for normality, which was found to be positively skewed for the 50 and 100 ms target durations, and to be leptokurtic for the 100 ms duration. The non-normality of these two conditions was confirmed through Shapiro-Wilk tests (all *p*s < .001), and for this reason non-parametric alternatives were chosen for the analysis.

A Kruskal-Wallis test showed the differences in congruency-related scene categorisation accuracy to be significantly affected by the presentation duration of target scenes, *H*(2) = 13.40, *p* = .001. Further to this, pairwise comparisons with Bonferroni-adjusted p values showed the disparity in performance differed significantly across target durations. The disparity in performance across congruency conditions at a target duration of 50 ms was significantly different to that at 100 ms (*p* = .004, *r* = 0.42), and at 250 ms (*p* = .005, *r* = 0.41). No significant difference was found between the performance disparity at 100 ms and 250 ms. Follow-up one-sample Wilcoxon tests were then used, comparing congruency-related differences in categorisation accuracy at each target duration to zero (representing no difference). Bonferroni adjusted p values are reported. At 50 ms, participants were significantly more accurate at categorising Congruous trials (*Mdn* = 0.87) than Incongruous trials (*Mdn* = 0.74), *Z* = −4.17, *p* < .001, *r* = −0.55, representing a large effect. The same pattern was true at 100 ms, with greater accuracy for Congruous trials (*Mdn* = 0.90) than Incongruous trials (*Mdn* = 0.87), *Z* = −2.68, *p* = .022, *r* = −0.35, representing a medium effect. This was also apparent at 250 ms, with greater accuracy for Congruous trials (*Mdn* = 0.97) than Incongruous trials (*Mdn* = 0.92), *Z* = − 3.27, *p* = .003, *r* = −0.44. This again represents a medium effect.

As predicted, the results from Experiment 1b mirrored those from Experiment 1a. Again, categorisation ability was significantly higher when target scenes were preceded by congruous approach images, and this differential in performance was greatest when target presentation duration was at its most brief (50 ms). Consequently, Experiment 1b confirmed our findings were due to the congruency manipulation, as opposed to simply being based on the presentation frequency of experimental trials. In addition, these results show that context-based predictions were being generated automatically by participants as they viewed the approach images, in line with work demonstrating that pre-target natural scene images lead to the automatic generation of expectations as to the identity of an upcoming target object (Caplette et al., 2020).

Taken together, the findings from across these two experiments revealed that approach images influenced subsequent scene processing, and so suggest a role for top-down information in rapid gist processing. They do not support, therefore, narratives which propose the extraction of scene-gist is exclusively based on feedforward processes.

## Experiment 2

While Experiments 1a-b found an influence of trial congruity, it remained to be determined the specific mechanisms responsible for such an advantage. Divergent explanations as to the mechanisms underlying the findings of the previous experiments are possible. On one hand, the presentation order of approach images may have comparatively little bearing on performance, whereby these images simply serve to provide a semantic context which increases the predictability of the subsequent destination. For instance, observing an approach image which depicts surroundings commonly associated with the countryside may be sufficient for expectations to be formed as to the most likely eventual destination (e.g., a field, woods, etc.). In this scenario there would be no cost to performance if approach images were not arranged in a meaningful sequence, as participants would still be provided with the same contextual information prior to target presentation. On the other hand, there may be an additional benefit, above-and-beyond that based on semantic context, from the spatiotemporally progressive nature of the series. If this were true, then we would expect to see lower performance on trials where there was disruption to the ordering of images within a sequence.

An advantage of approach-image sequentiality, if apparent, could be due to several factors. For example, the importance of narrative coherence for efficient processing has previously been demonstrated (Cohn et al., 2012; Smith & Loschky, 2019). Through disruption to the order and content of comic strips, Cohn and colleagues have investigated the individual contributions of the semantic relationship between images and their overall narrative structure on the processing of image sequences (2012). It was found that both semantic relatedness and narrative structure were advantageous, whereby the processing of a subsequent image was influenced by both the structure and meaning of the series that preceded it. Alternatively, a case could be made that sequentiality allows for the generation of a ‘perceived flow’ of movement through the environment, potentially facilitating processing by allowing for the extraction of more information, such as that derived from the semblance of optical flow (Gibson, 1966) or through aiding the transformation of the viewer-centred 2½D sketch into a three-dimensional representation (Marr, 1982). Finally, the further away in space a leading image is from its eventual destination, the potentially weaker its predictive power. As an observer progresses through a series, each new leading image may further ‘fine-tune’ expectations, which could be a more additive process compared to that occurring from experiencing the same images in random order.

Hence, to investigate whether the sequentiality of series plays a role in gist processing, in Experiment 2 we manipulated the presentation order of approach images while also continuing to manipulate congruency. We predicted a categorisation advantage for sequentially coherent trials, as compared to disordered trials. This was due to the assumption that sequentiality would create a flow of information that more closely mirrored typical functioning in everyday environments, and due to research identifying an important role of narrative sequences for processing (e.g., Cohn et al., 2012; Smith & Loschky, 2019). Additionally, owing to the provision of semantically relevant context, we predicted that performance on Congruous trials, regardless of sequentiality, would still exceed that on Incongruous trials.

### Design

While maintaining the approach-destination congruency manipulation of Experiments 1a-b, Experiment 2 departed from the previous iterations by also manipulating the sequentiality of approach images. Therefore, trials included approach images displayed either in a sequential or randomised order. This led to four within-participant conditions: Congruous-Sequential; Congruous-Disordered; Incongruous-Sequential; and Incongruous-Disordered. Each condition consisted of 30 trials and included one series for each of the scene categories. A Latin Square design was employed, so that each series alternated across all conditions within the four versions of the experiment. For each version, the destination images for those series selected to constitute Incongruous trials were randomly reallocated amongst each other, following the same principles as previous iterations. Similarly, the presentation order of approach images within Disordered trials was randomly selected, but with two important constraints. First, the approach image in the closest geographical location to the destination scene could not be the final pre-target image in a Disordered trial. This parameter was to ensure that congruous targets were not simply being primed by the final approach image in isolation. Secondly, such trials could not contain more than two approach images displayed in their original order. This was to safeguard the non-sequentiality of Disordered trials.

The presentation order of trials was randomised independently for each participant. Target duration was not manipulated in Experiment 2. Targets were presented for 50 ms, due to the findings from the previous experiments. This was based on the demonstration that the effect of congruity was most apparent at brief target durations, diminishing as presentation length increased. See Figure 4 for a schematic of the experimental protocol.

**Figure 4.**
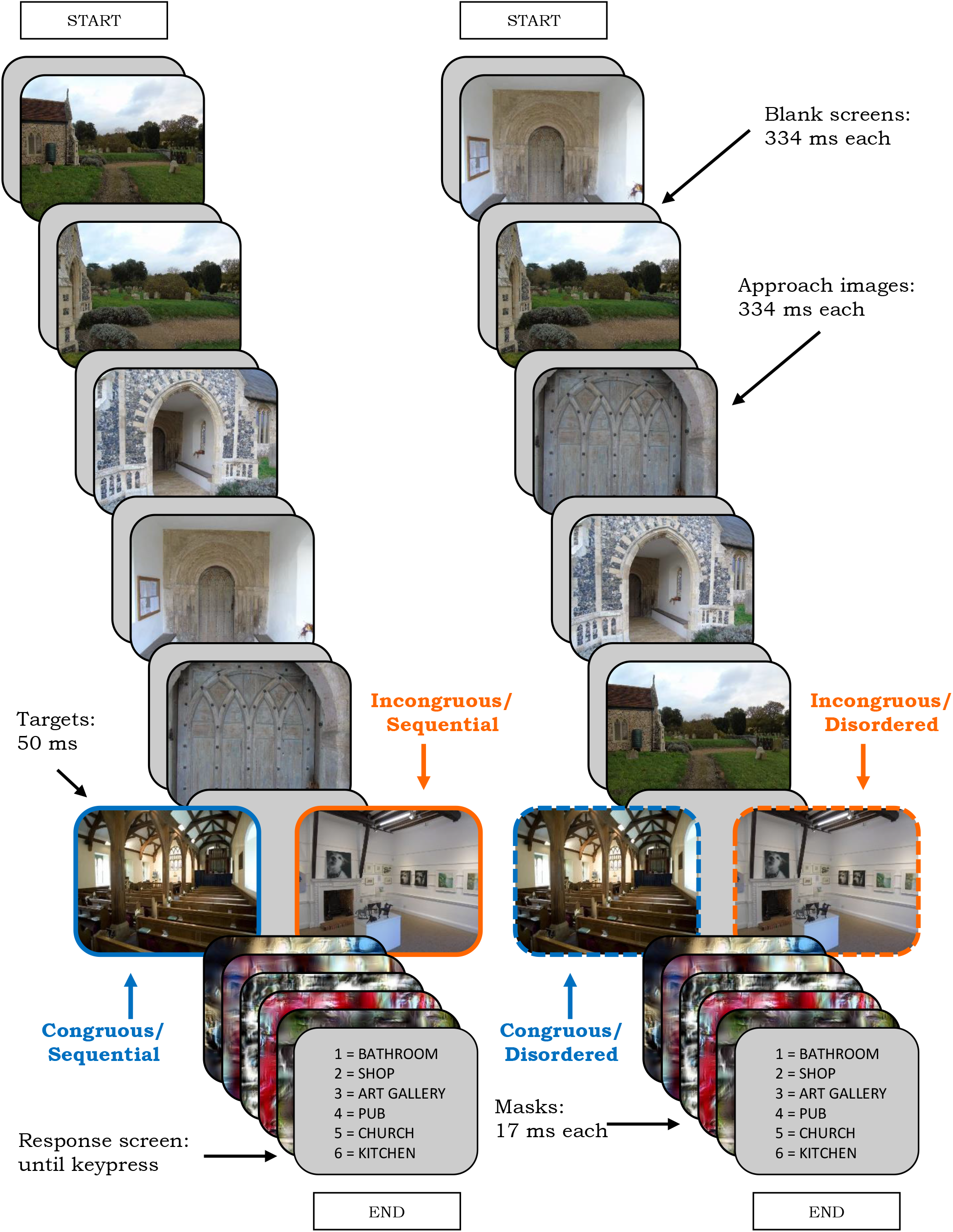
Schematic of the Protocol for Experiment 2

### Participants

Thirty-six participants were originally included in Experiment 2, made up from both students and staff of the university, receiving either course credits or a small payment for taking part. However, analysis of this initial data showed a much smaller effect of approach-image sequentiality compared to the size of effect related to trial congruency. To better determine the veracity of this effect, we took the decision to double the sample size while halving the alpha level during the subsequent analysis (*α* = .025). This technique is considered appropriate for controlling the Type 1 error rate in situations where a sample is increased due to the size of the observed effect (Lakens, 2014). The complete sample, therefore, consisted of 72 participants (*M*_age_ = 23.58, *SD*_age_ = 11.22; 54 Females, 18 Males; 61 Right-handed, 10 Left-handed, 1 Ambidextrous).

### Stimuli

Experiment 2 used the same image set and masks as Experiment 1a and 1b. The response screens were again redrawn, using the same randomisation procedures as the preceding experiments.

### Procedure

The procedure followed the same routine as Experiment 1b, except that the display duration of blank screens was increased from 167 ms to 334 ms (10 to 20 frames on a 60Hz monitor). This was judged to provide a more comfortable viewing experience for participants, which better mimicked the sense of traversing an environment.

## Results and Discussion

One participant was removed due to a score outside three standard deviations of the mean in one of the experimental conditions. Analysis was conducted on the remaining 71 participants (*M*_age_ = 23.62, *SD*_age_ = 11.29; 54 Females, 17 Males; 60 Right-handed, 10 Left-handed, 1 Ambidextrous). For each participant, accuracy scores for Congruous-Disordered trials were subtracted from scores for Congruous-Sequential trials, and Incongruous-Disordered scores were subtracted from Incongruous-Sequential scores, to assess performance differences across Congruency and Sequentiality conditions. Additionally, differences across Incongruous trials were subtracted from the differences across Congruous trials. These three sets of values were used to test the data for normality, displaying no issues relating to skewness or kurtosis, as confirmed through Shapiro-Wilk tests (all *p*s > .3). On account of this, a 2 (Congruency: Congruous; Incongruous) x 2 (Sequentiality: Sequential; Disordered) repeated measures ANOVA with planned comparisons was chosen for the analysis.

There was a main effect of Congruency, *F*(1, 70) = 9.74, *p* = .003, ƞp^2^ = .12, with significantly higher performance on Congruous (*M* = 0.77, *SE* = 0.02) than Incongruous (*M* = 0.69, *SE* = 0.03) trials. This represents a medium effect size. There was no main effect of Sequentiality, *F*(1, 70) = 0.32, *p* = .574, ƞp^2^ = .01. There was, however, a significant Congruency X Sequentiality interaction, *F*(1, 70) = 7.00, *p* = .010, ƞp^2^ = .09, also representing a medium effect size. This indicated that the sequentiality of approach images had different effects on categorisation performance depending on whether the approach images were congruous with the target image (see Figure 5). To investigate this interaction further, paired samples t-tests were conducted. On average, when approaches were congruous with destinations, participants performed better if the approach images were presented in sequential (*M* = 0.79, *SE* = 0.02) compared to random order (*M* = 0.76, *SE* = 0.02). This difference, 0.03, 95% CI [0.003, 0.05], was not significant at the 0.025 alpha level, *t*(70) = 2.24, *p* = .028, *r* = .26. This represents a small-to-medium effect. It should be noted, though, that the confidence interval did not bridge zero which can be taken as support for the existence of such an effect. When approaches were incongruous with destinations, participants scored slightly higher if approach images were presented in random order (*M* = 0.69, *SE* = 0.03) compared to sequentially (*M* = 0.68, *SE* = 0.03). However, this difference, −0.02, 95% CI [-0.05, 0.01] did not reflect a significant difference in performance, *t*(70) = −1.16, *p* = .249, *r* = .14.

**Figure 5.**
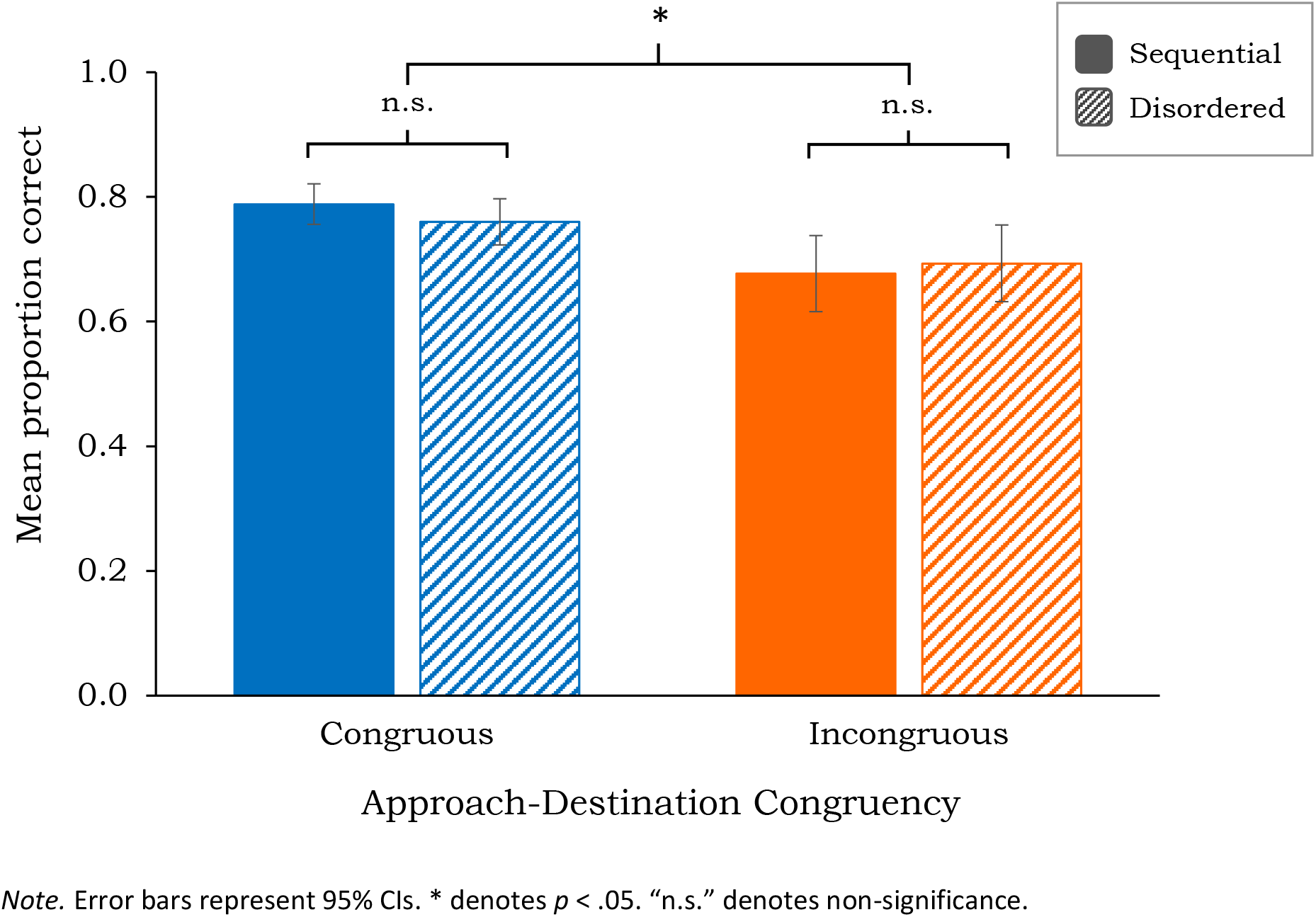
Scene Categorisation Accuracy for Sequential and Disordered Trials as a Function of Approach-Destination Congruency

As with the previous experiments, we again found a benefit to categorisation performance related to trial congruency. However, the predicted effect of approach-image sequentiality did not reach statistical significance. As a consequence, it appears that simply providing observers with an appropriate contextual setting allowed for sufficiently accurate predictions of the upcoming destination to be formed. For instance, approach images displaying movement along a pavement, even if out of sequence, still allowed for expectations to be generated (i.e., previous experience would teach us that pavements lead to, say, high streets much more frequently than to woods).

That there was no significant additional benefit to gist processing when trials depicted a continuous spatiotemporal journey suggests that the generation of a perceived ‘flow of movement’ (e.g., Gibson, 1966) had little bearing on the accuracy of the expectations constructed by participants. This finding was also surprising due to research demonstrating the importance of narrative coherence within pre-target sequences (Cohn et al., 2012; Smith & Loschky, 2019). However, that previous work incorporated narratives more complex in nature than our short approaches to proximal destinations. It stands to reason that the disruption to processing caused by the disarrangement of chronological elements would be a function of the complexity of the story being told, whether that be within the panels of a comic strip or the pictorial representation of an extended journey.

## Experiment 3

The findings from Experiment 2 suggested that the influence of approach images on subsequent scene processing was primarily due to participants being provided a semantic context prior to target onset. Up to this point the assumption had been made that this effect was driven by congruous approaches facilitating the subsequent processing of destinations. However, a possible alternative explanation remained, namely that the difference across experimental conditions was the result of incongruous approaches interfering with the processing of destinations.

To answer this question, in Experiment 3 we introduced a third, neutral condition whereby approach images were replaced by images of coloured patterns. As such, provision of semantic context was absent within the trials of this condition, meaning participants were unable to generate expectations as to the identity of the upcoming destination. As this condition maintained the trial format of other conditions, namely the display of series of pre-target images, it was considered a suitable reflection of baseline performance. Therefore, by comparing categorisation performance for this condition to performance on Congruous and Incongruous trials, respectively, we hoped to uncover more fully the role of the congruency manipulation on gist processing. We expected better performance on Congruous trials, compared to No-context and Incongruous trials; due to a lack of direct evidence from previous research, we made no predictions as to whether performance on Incongruous trials would be significantly lower than that of the No-context condition.

### Design

Experiment 3 maintained the approach-destination congruency manipulation of previous experiments but did not include the manipulation of approach sequentiality seen in Experiment 2. Alongside the previous congruency conditions, a third condition was added in which the approach images provided no semantic context to participants prior to destination-onset. This led to three within-participant conditions: Congruous; Incongruous; and No-context. Each condition consisted of 40 trials, including at least one, and no more than two, series for each of the scene categories. A Latin Square design was employed, so that each series alternated across all conditions within the three versions of the experiment. For each version, the destination images for those series selected to constitute Incongruous trials were randomly reallocated amongst each other, in line with the principles of previous iterations. The presentation order of trials was randomised independently for each participant. Target duration was again not manipulated in Experiment 3, as the presentation of targets was set at 50 ms for all participants. See Figure 6 for a schematic of the experimental protocol.

**Figure 6.**
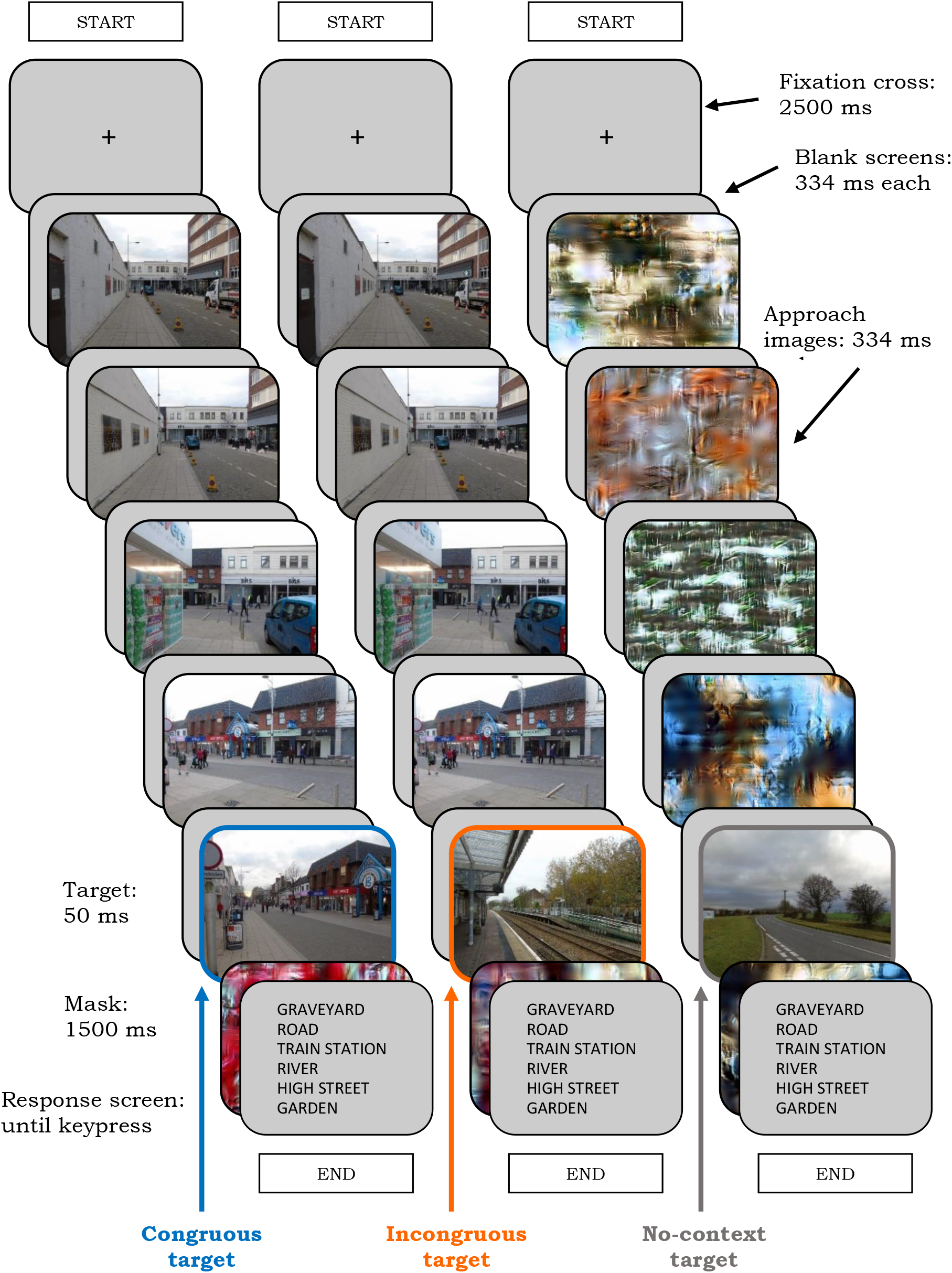
Schematic of the Protocol for Experiment 3

### Participants

Participants were recruited through the online international participant pool, Prolific (www.prolific.co), and received a small payment for taking part. Demographic screeners were used to ensure all participants were adults who lived in the UK, US, Canada, Australia or New Zealand, and were fluent speakers of English. This filtering was to ensure that all participants would both be able to fully understand the task instructions and would be familiar with the types of sceneries used in the experiment. Forty-five participants took part in Experiment 3 (*M*_age_ = 33.53, *SD*_age_ = 11.62; 30 Females, 15 Males; 37 Right-handed, 7 Left-handed, 1 Ambidextrous).

### Stimuli

Experiment 3 used the same image set and masks as the previous experiments, although in this iteration some of the mask-images were repurposed to act as leading images in the No-context condition (as set out below). The response screens were again reconfigured, using the same randomisation procedures as before.

### Procedure

The procedure followed a similar routine as Experiment 2, with some minor alterations necessary for the experiment to be run online. The experiment was programmed using Testable (www.Testable.org) and, due to constraints imposed by the software, the number of images displayed per trial needed to be reduced. This was achieved in two ways. Firstly, only four approach images were presented per trial, as we removed the first approach image from each series (i.e., the image most geographically distant from the destination). Secondly, destination images were followed by a single mask rather than a set of five dynamic masks. The duration of these individual masks was extended to 1500 ms to ensure a suitable disruption to processing from target-offset was maintained. The previous experiments each used 600 mask-images, and so 120 of these were randomly selected to again be used as masks in Experiment 3. A further 160 were then randomly selected to serve as leading images in the No-context condition. The order of presentation of these images, both within series and across trials, was also randomised. These randomisation procedures were followed for each of the three Latin Square versions, with the proviso that a mask-image could not be used as a leading image and a mask within a single version, and that all 600 mask-images were used across the experiment as a whole.

Two further minor alterations were included to ensure the smooth running of the experiment, due to the inevitable reduction in researcher oversight during an online study. Firstly, a fixation cross was displayed in the centre of the screen prior to the start of each series. Secondly, selection of a response was made by navigating a cursor to the chosen textbox, rather than by pressing a number on a keypad.

## Results and Discussion

All participants scored within three standard deviations of the mean for each of the experimental conditions, and so all 45 were included in the analysis. For each participant, the difference in accuracy scores for the Congruous compared to the No-context condition, and the No-context compared to the Incongruous condition, were calculated. These values revealed the data to be normally distributed, displaying no issues with skewness or kurtosis, as confirmed through Shapiro-Wilk tests (all *p*s > .3). Consequently, a one-way (Congruency: Congruous; Incongruous; No-context) repeated measures ANOVA with planned comparisons was chosen for the analysis.

Mauchly’s test indicated that the assumption of sphericity had been violated, *X*^2^(2) = 16.00, *p* < .001, and so degrees of freedom were corrected using Greenhouse-Geisser estimates of sphericity (Greenhouse & Geisser, 1959). There was a significant effect of the type of approach image on categorisation performance, *F*(1.53, 67.18) = 78.37, *p* < .001, ƞp^2^ = .64. This represents a large effect. As can be seen in Figure 7, the proportion of correct responses was greatest in the Congruous condition (*M* = 0.79, *SE* = 0.01), followed by the No-context condition (*M* = 0.70, *SE* = 0.02), and with weakest performance in the Incongruous condition (*M* = 0.52, *SE* = 0.03). Follow-up paired samples t-tests, with Bonferroni-adjusted p values, revealed that the mean performance difference between Congruous and No-Context trials, 0.10, 95% CI [0.07, 0.13], was significant *t*(44) = 6.21, *p* < .001, *r* = .68, as was the difference between Congruous and Incongruous trials, 0.27, 95% CI [0.22, 0.32], *t*(44) = 10.19, *p* < .001, *r* = .84. Both represent large effects. Furthermore, the mean performance difference between No-context and Incongruous trials, 0.17, 95% CI [0.13, 0.22], was also found to be significant *t*(44) = 7.84, *p* < .001, *r* = .76, again representing a large effect.

**Figure 7.**
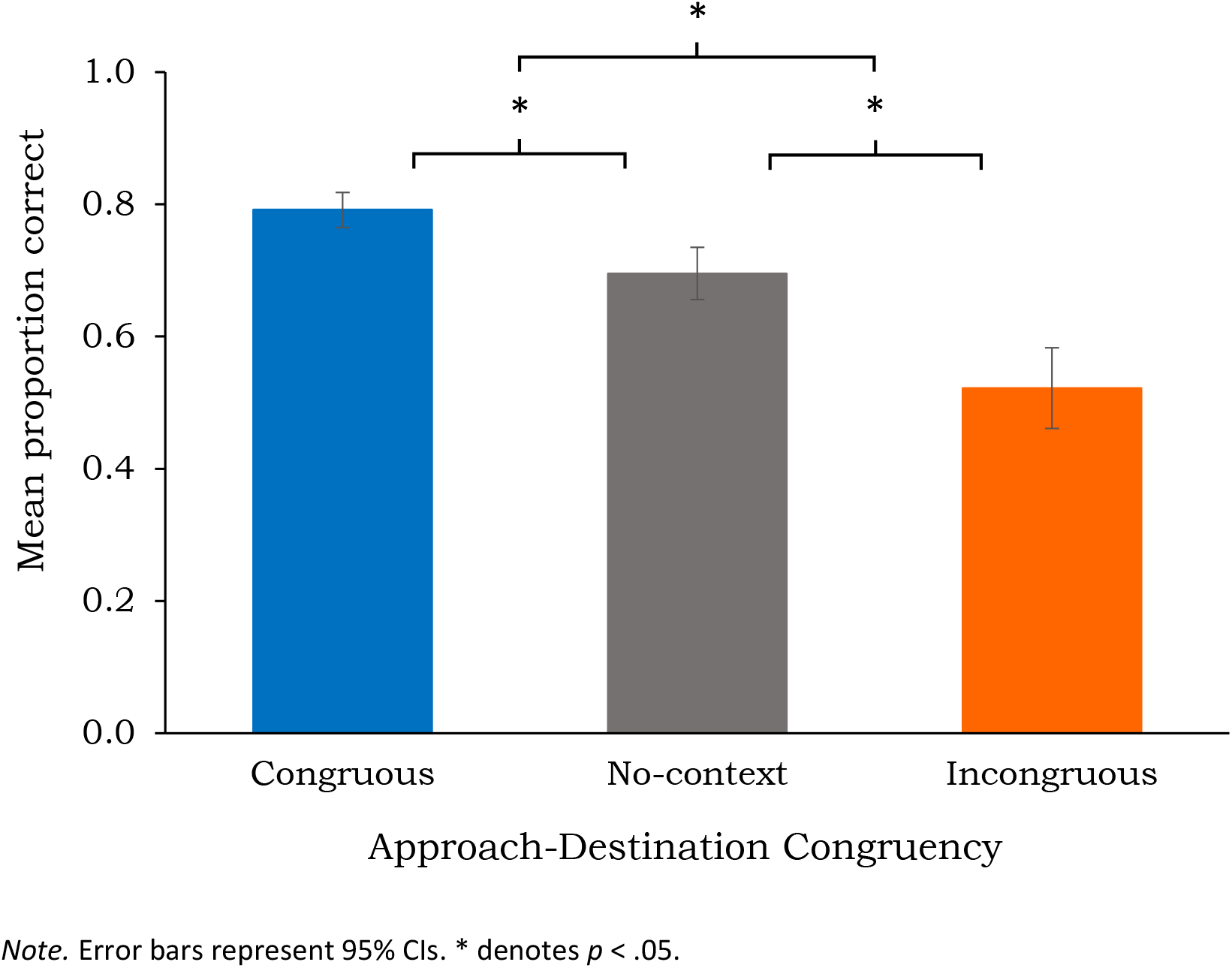
Scene Categorisation Accuracy for Congruous, Incongruous and No-context Trials

A clear pattern of results emerged in Experiment 3, with significantly contrasting levels of categorisation performance seen across each of the three conditions. As with previous iterations, the ability to categorise destination scenes was greater when preceded by semantically congruous, rather than incongruous, approaches. Further to this, increased performance was also apparent on Congruous trials as compared to those where pre-target context was absent, thus confirming our contention that semantic congruity leads to a facilitation of gist processing.

In addition, we found that performance on Incongruous trials was significantly below that of the No-context condition, revealing that participants’ ability to categorise a destination scene was inhibited if preceded by an unrelated scenic context. This appears to be in agreement with previous research showing scenes containing unexpected features are more difficult to extract meaning from (Greene et al., 2015), as well as recent work demonstrating interference to object recognition as a result of contextual violations within object-scene pairs (Lauer et al., 2020). In sum, this pattern of results clearly shows that approach images were eliciting expectations as to the likely identity of an upcoming target scene, resulting in a benefit to gist processing if expectations were realised but, alternatively, resulting in a cost to processing if violated.

## Experiment 4

The initial set of behavioural experiments revealed that providing semantic information leads to increased performance on subsequent scene processing. Building on these results, we turned to an investigation of the neural signature. To that end, in Experiment 4 we investigated the event-related potential (ERP) correlates of scene processing and the role of expectations on gist extraction. This investigation was exploratory in nature as, to date, we are unaware of any prior use of this methodology for the examination of the role of sequential, naturalistic leading images on subsequent scene-gist processing. However, as set out below, previous research allowed for inferences to be made as to the ERP components most likely to be correlated with the effect of congruency on scene processing.

Using such a methodology provides the opportunity to better understand the timing of expectation-related alterations to gist processing. While manipulation of target presentation duration in behavioural studies can help point towards the general speed of an effect, it alone cannot determine the point of occurrence for expectation-induced violations to scene processing; despite utilising dynamic backward masking, a complete cessation to target-image processing at offset is unlikely. Event-related potentials, on the other hand, offer a more precise means by which to uncover the temporal points at which differences in brain activity emerge as a function of scene congruency, allowing for conclusions to be drawn as to the potential swiftness of any top-down influence.

Therefore, the first ERP component selected for investigation was the P2. Arising rapidly within Parieto-occipital regions – at around 200 ms after target onset – this component has been proposed as the earliest known marker for scene-specific processing (Harel et al., 2016), affected by changes in global scene properties but not top-down observer-based goals (Hansen et al., 2018). However, the exact influence of top-down information on the P2 remains unclear. While evidence indicates early components such as this are sensitive to low-level visual information such as salience (Straube & Fahle, 2010), as well as object identification (Viggiano & Kutas, 2000), the influence of higher-level processes is less well determined. For example, differences in ERPs at around 200 ms have been found when identifying the presence of objects within briefly presented natural images, potentially reflecting decision-related activation (Thorpe et al., 1996; VanRullen & Thorpe, 2001). As a result, a lack of agreement exists, both in terms of whether the P2 is altered by top-down processing at all and, if so, what form of top-down processing might hold influence. Furthermore, it has been suggested the P2 may in fact index an intermediary processing stage, somewhat bridging perceptual and higher-order processes, such as segmentation and categorisation, respectively (De Cesarei et al., 2013). In terms of the current study, the above implies predictions relating to the P2 must be tentative. We can contend, however, that if congruency-based differences in activation were shown to exist within the earliest indicant of scene-specific processing (Harel et al., 2016), this would be representative of expectations influencing early perceptual processing. More broadly, finding such activation differences would signify that the P2 component is open to influence from top-down information. Indeed, top down modulation of the P2 as being related to the semantic processing of scenes has been proposed before (Federmeier & Kutas, 2002).

As well as determining the timing of initial alterations to gist processing, ERP analysis can help elucidate the mechanisms underlying expectation-related performance changes as cognitive processing continues. In other words, investigating activation changes across subsequent scene-related ERP components can help reveal the manner in which scene congruity might affect gist processing. Previous scene processing research has shown two later components as being susceptible to experimental manipulation. The first of these – the N400 – has long been associated with semantic processing, with its amplitude observed as being inversely proportional to semantic expectancy (e.g., Kutas & Hillyard, 1984) and more generally to the ease with which conceptual information can be retrieved (Van Petten & Luka, 2006). For this component a certain level of consensus has been reached: across both central and anterior sites, increased negativity within the N400 time window has been related to scene-object semantic violations in static images (Ganis & Kutas, 2003; Mudrik et al., 2010; Võ & Wolfe, 2013) and within video clips (Sitnikova et al., 2003; Sitnikova et al., 2008). N400 effects have also been found to be sensitive to the semantic association between pairs of sequential pictures (Barrett & Rugg, 1990), and to violations of semantic expectation in language comprehension studies (Holcomb, 1993; Van Petten, 1995). A similar pattern of N400 changes across conditions within the current study would, therefore, indicate that differential behavioural performance derived from congruency-based manipulations in the behavioural experiments was due to semantic violations, rather than simply violations of expected low-level visual information.

The second of these later components, again potentially revealing in terms of the mechanisms responsible for the effect of expectations on processing, is the P600. Like the N400, this component was initially described in language comprehension studies, where syntactic errors creating a need for sentence reanalysis were observed to elicit increased positivity within posterior regions at ∼600 ms (Hagoort et al., 1993; Osterhout & Holcomb, 1992). This was irrespective of whether sentences were experienced through visual or auditory modalities (Hagoort & Brown, 2000; Osterhout & Holcomb, 1993). Similarly, within scene processing research, increased positivity at the P600 has been reported as reflecting reanalysis prompted by mis-located objects (Võ & Wolfe, 2013). There, increased late positivity was found when appropriate objects were positioned in inappropriate places within a scene (such as a dishtowel on a kitchen floor), irregularities proposed by the authors as reflecting syntactic – rather than semantic – violations. However, a lack of agreement should be noted regarding the functional role of the P600. For instance, It has been suggested that this increased late positivity may not exclusively represent syntactic violations, as its sensitivity to semantic information has also been demonstrated (Gunter et al., 1997; Gunter et al., 2000; Kuperberg, 2007; Sitnikova et al., 2003).

Furthermore, such changes to late positive components have not always been observed when objects break syntactic rules within scenes (e.g., Demiral et al., 2012). To confuse matters further, while Võ and Wolfe (2013) did not find alterations to the P600 when inappropriate objects were placed in appropriate locations – taken by the authors as evidence of the dissociation between the effects of semantic and syntactic violations – this form of semantic violation was shown to elicit a reduction in P600 amplitude in previous work (Mudrik et al., 2010). It is possible this inconsistency across studies is rooted in contrasting methodological choices, with one allowing for expectations to be generated due to the context-scene appearing prior to the target object (Võ & Wolfe, 2013), and the other avoiding this through simultaneous presentation of targets and their associated scenes (Mudrik et al., 2010).

There is still much debate as to the comparability between language and scene processing in general, particularly in terms of whether the processing of words and pictures shares a common semantic system (e.g., Federmeier & Kutas, 2002), and questions remain for both paradigms in relation to the nature of the P600. However, perhaps the most reproducible findings regarding this component have been through the use of ‘garden path’ sentences in linguistic studies, whereby violation of the expected structure of a sentence creates the need for reanalysis of the preceding sequence of words (e.g., Frazier & Fodor, 1978; Osterhout & Holcomb, 1992). Accordingly, display of similar congruency-related changes to the P600 in the current study would suggest the violation of expectations, created through approach images, resulted in the need for reanalysis of incongruous targets. In other words, just as a garden path sentence might build an inaccurate expectation as to the grammatical structure of a sequence of words, which is subsequently violated, a sequence of images depicting a journey is likely to build expectations as to the eventual destination, only for this to be violated on presentation of an incongruous target scene.

Based on the above, predictions as to the pattern of results can be made, although they must remain speculative due a lack of consensus across previous research. Firstly, the association between the N400 and semantic incongruity, observed in studies of language and scene processing, leads us to expect greater N400 amplitudes across central and anterior regions for Incongruous trials, as compared to Congruous trials. Secondly, due to violations in thematic coherence, we expect to observe increased P600 amplitude across posterior sites for Incongruous trials, as compared to Congruous trials. Finally, due to debate remaining as to the influence of top-down factors on the P2 component, we do not make predictions as to whether Incongruous trials will elicit increased positivity in posterior regions during this time-window. However, if such changes were observed, we would take this as signalling the violation of expectations was able to influence the earliest stages of scene processing, including the integration of visual properties.

### Design

To maintain consistency throughout the study the experimental protocol mirrored previous iterations closely, although with certain alterations necessary to improve the suitability of the trial routine for use with electroencephalography. The 120 experimental trials were split equally across two conditions of approach-destination congruity, and there was no manipulation of approach sequentiality. Sixty image-series were randomly selected to serve as Incongruous trials, with their destination images randomly redistributed amongst themselves. The same restrictions were applied to the randomisation procedure as previous experiments. A counterbalanced version of the protocol was created, and these two versions were employed in a Latin Square across the course of the experiment to ensure no unintended bias was introduced due to the allocation of trials to congruency conditions. See Figure 8 for a schematic of the experimental protocol.

**Figure 8.**
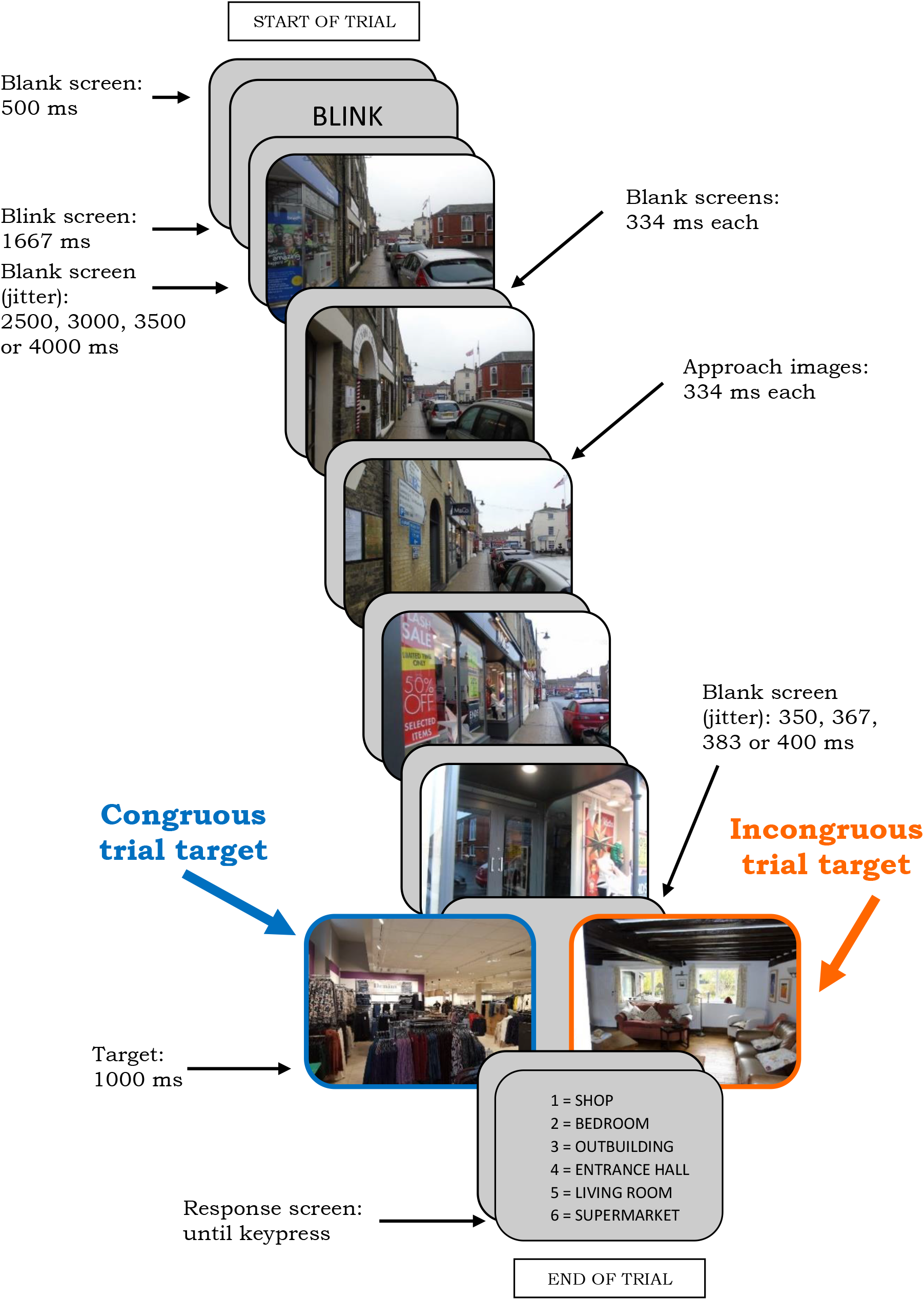
Schematic of the Protocol for Experiment 4

### Participants

Experiment 4 included 26 Psychology students, again recruited through the research pool and given course-related credits for taking part (*M*_age_ = 20.31, *SD*_age_ = 2.77; 20 Females, 6 Males; 19 Right-handed, 7 Left-handed). None had participated in any of the behavioural experiments, and so all were unfamiliar with the stimuli and naïve to the purpose of the study. One participant was removed as their comprehension of the task could not be assured (incorrectly responding to 77% of Incongruous trials), and another removed due to excessive high-frequency noise across multiple channels. Analyses were conducted on the remaining 24 participants (*M*_age_ = 20.42, *SD*_age_ = 2.86; 18 Females, 6 Males; 18 Right-handed, 6 Left-handed). All participants reported as having no history of neurological disorders.

### Stimuli

The same image set was again used, although all masks were removed for Experiment 4. Response screens were reconstructed using the same guidelines as previous experiments.

### Procedure

Each trial began with a ‘blink’ screen, followed by a blank screen including a jitter (duration: 2.5, 3, 3.5 or 4 seconds). This was to protect the ERPs from the potential systematic influence of slow baseline drifts coinciding with the routine. The jitter was pseudo-randomised to ensure a different blank screen duration prior to each of the four approach-series per scene category. These initial screens gave participants the opportunity to get comfortable prior to the presentation of each trial, with the aim of reducing the number of movement-based artefacts within the subsequent ERPs. A second, shorter jitter (duration: 350, 367, 383 or 400 ms) was also introduced to the last blank screen prior to target presentation, to shield against artefacts caused by participants being able to predict the exact onset time of the target. This jitter was pseudo-randomised in the same manner as before, and was evenly distributed across the two congruency conditions. There was no manipulation of target duration, with presentation length set at 1 second. This extended duration served two purposes: firstly, as only correctly answered trials were used in the analysis it sustained a high level of categorisation performance and, secondly, it protected against noise within the ERP caused by the offset of the stimulus or the onset of the response screen. No masking was used, with the target followed by a blank screen prior to a 6AFC response screen.

### Data Acquisition

The EEG was recorded using a Brain Vision 64-channel active electrode system, embedded within a nylon cap (10/20 system). Electrode FT9 was removed from the cap and placed under the left eye to monitor blinks and eye movements. The signal was acquired at a 1000 Hz sampling rate with FCz used as the online reference (see Figure 9).

**Figure 9.**
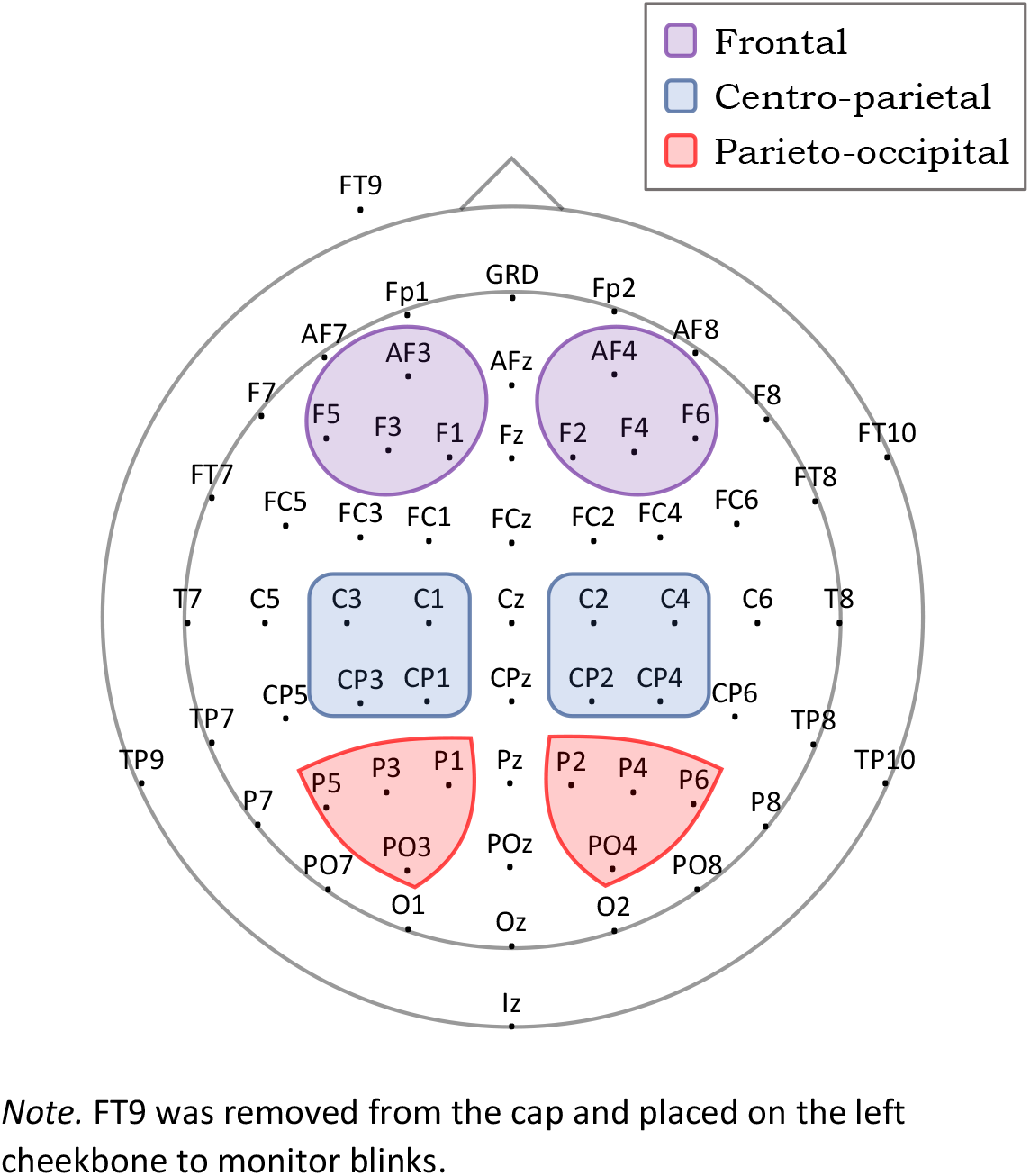
Map of Electrode Placement Including the ROIs

### Processing

Offline processing and analyses were conducted using EEGLAB (Delorme & Makeig, 2004) and ERPLAB (Lopez-Calderon & Luck, 2014), running under Matlab 9.2.0 (R2017a, Mathworks, Natick, MA). Trials with incorrect responses were removed from the continuous EEG (3.89% of Congruous trials and 5.01% of Incongruous trials across participants). Ocular artefact correction took place through Independent Component Analysis (ICA) to identify blinks and lateral eye movements. These artefacts are located at anterior electrodes and can be identified based on their characteristic shapes (frequent clear spikes or step-like functions, respectively). Therefore, removal of these components was conducted manually by simultaneously comparing the continuous EEG to the time-course of the Independent Components. This led to removal of 41 Independent Components across the sample as a whole, with no more than two components removed for any single participant. Re-referencing to the average of the TP9 and TP10 electrodes (which approximate to the location of the mastoids) was computed offline (e.g. Cohn & Foulsham, 2020). Any channels suffering from persistent high-frequency noise were interpolated using the mean signal from the surrounding electrodes (mean percentage of channels interpolated across participants: < 1%). After removal of DC trends, an IIR Butterworth filter was applied for high- and low-pass filtering the data with half-amplitude cut off values of 0.01 Hz and 80 Hz, respectively (12 dB/oct; 40 dB/dec). The EEG was segmented into epochs of 1 second, from 200 ms before to 800 ms after target-scene onset. The length of the baseline used to correct epochs was the 200 ms immediately preceding target onset. Epochs contaminated with excessive artefacts were identified, and rejected, by setting a peak-to-peak voltage threshold of 100 µV across a moving window of 200 ms with a window step of 50 ms. This resulted in the rejection of 6.94% of Congruous trials and 7.10% of Incongruous trials across participants.

The amplitudes of the P2, N400 and P600 were measured as the mean of all data points between 175-250 ms, 300-500 ms and 500-700 ms, respectively. These specific components were chosen as the P2 has previously been suggested as the earliest indicator of scene selectivity (Harel et al., 2016), while the N400 and P600 have been associated with semantic and syntactic integration, respectively (e.g., Friederici et al., 1993; Hagoort & Brown, 2000; Holcomb, 1993; Mudrik et al., 2010; Van Petten, 1995; Võ & Wolfe, 2013).

The time windows chosen are commonly used as boundaries for investigating the N400 (e.g., Ganis & Kutas, 2003; Guillaume et al., 2016; Mudrik et al., 2010) and P600 (Angrilli et al., 2002; Cohn et al., 2014; De Vincenzi, 2003). Less standardisation exists regarding the P2, however, with previous research involving the processing of scenes employing time windows ranging anywhere between 140 to 320 ms post-stimulus onset (see, for example, De Cesarei et al., 2013; Ferrari et al., 2017; Harel et al., 2020; Yuan et al., 2007). We, therefore, determined our window of interest based on visual inspection of the grand average ERP. As a result, a window of 175-250 ms was selected as it covered the 220 ms timepoint previously identified as showing maximal amplitude for scene processing (Harel et al., 2016), while offering as large a span as was achievable without incorporating elements of the proximal P1 and P3 components.

Key electrode sites were grouped into three regions of interest (ROIs), each incorporating eight electrodes (split equally across hemispheres). A Centro-parietal ROI included electrodes C1/C2, C3/C4, CP1/CP2 and CP3/CP4, a Parieto-occipital ROI comprised electrodes P1/P2, P3/P4, P5/P6 and PO3/PO4, and a Frontal ROI contained electrodes F1/F2, F3/F4, F5/F6, and AF3/AF4 (see Figure 9). The posterior ROI was selected as Parieto-occipital regions are associated with maximal amplitude of the P600 (e.g., Gouvea et al., 2010) and the P2 (e.g., Hansen et al., 2018). The more central and anterior ROIs were chosen as the amplitude of the N400 has previously been found to be maximal at Centro-parietal regions (e.g., Ganis & Kutas, 2003), while the processing of semantic information related to images, as compared to text, has often been shown to elicit a Frontal negativity during the 300-500 ms temporal window (e.g., Ganis et al., 1996; Holcomb & McPherson, 1994; Mudrik et al., 2014).

## Results

Analysis was conducted on the mean amplitudes for each time-period of interest using 2 (Hemisphere: Left; Right) x 3 (Region: Centro-parietal; Parieto-occipital; Frontal) x 2 (Congruency: Congruous; Incongruous) repeated-measures ANOVAs. Where Mauchly’s test revealed possible violations of the sphericity assumption Greenhouse-Geisser corrected values are reported (Greenhouse & Geisser, 1959). Significant interactions were followed up with paired t-tests where appropriate. See Appendix C for a summary of the statistical analyses conducted.

### 175-250 ms Window

A three-way ANOVA revealed no main effect of Congruency (*p* = .842). There was also no three-way interaction (*p* = .552), nor a Hemisphere x Region interaction (*p* = .396), nor a Hemisphere x Congruency interaction (*p* = .424). There was, however, a significant Region x Congruency interaction *F*(2, 46) = 15.68, *p* < .001, ƞp^2^ = .41. See Figure 10 for scalp maps of voltage differences across conditions. In terms of this interaction, follow-up paired t-tests revealed no significant effect of congruency within the Centro-parietal ROI (*p* = .893). There was a significant effect within the Frontal region, *t*(23) = 2.54, *p* = .018, *r* = .47, due to there being a significantly more negative mean amplitude for Incongruous trials (*M* = −3.06) than Congruous trials (*M* = −2.27). See Figure 11 for the grand-averaged Frontal ERPs. This represents a medium-to-large effect. There was also a significant effect within the Parieto-occipital region, *t*(23) = −2.08, *p* = .048, *r* = .40, representing a medium-sized effect. This was due to there being a significantly more positive mean amplitude for Incongruous trials (*M* = 4.90 µV) than Congruous trials (*M* = 4.33 µV) within Parieto-occipital areas (see Figure 12 for the grand-averaged Parieto-occipital ERPs). Additionally, we re-ran our analysis at slightly more lateral posterior sites, in regions where maximal P2 changes have previously been shown (e.g., Harel et al., 2016; Harel et al., 2020; Hansen et al., 2018). This confirmed our finding of congruency-related changes to the P2 component (see Appendix D for further details).

**Figure 10.**
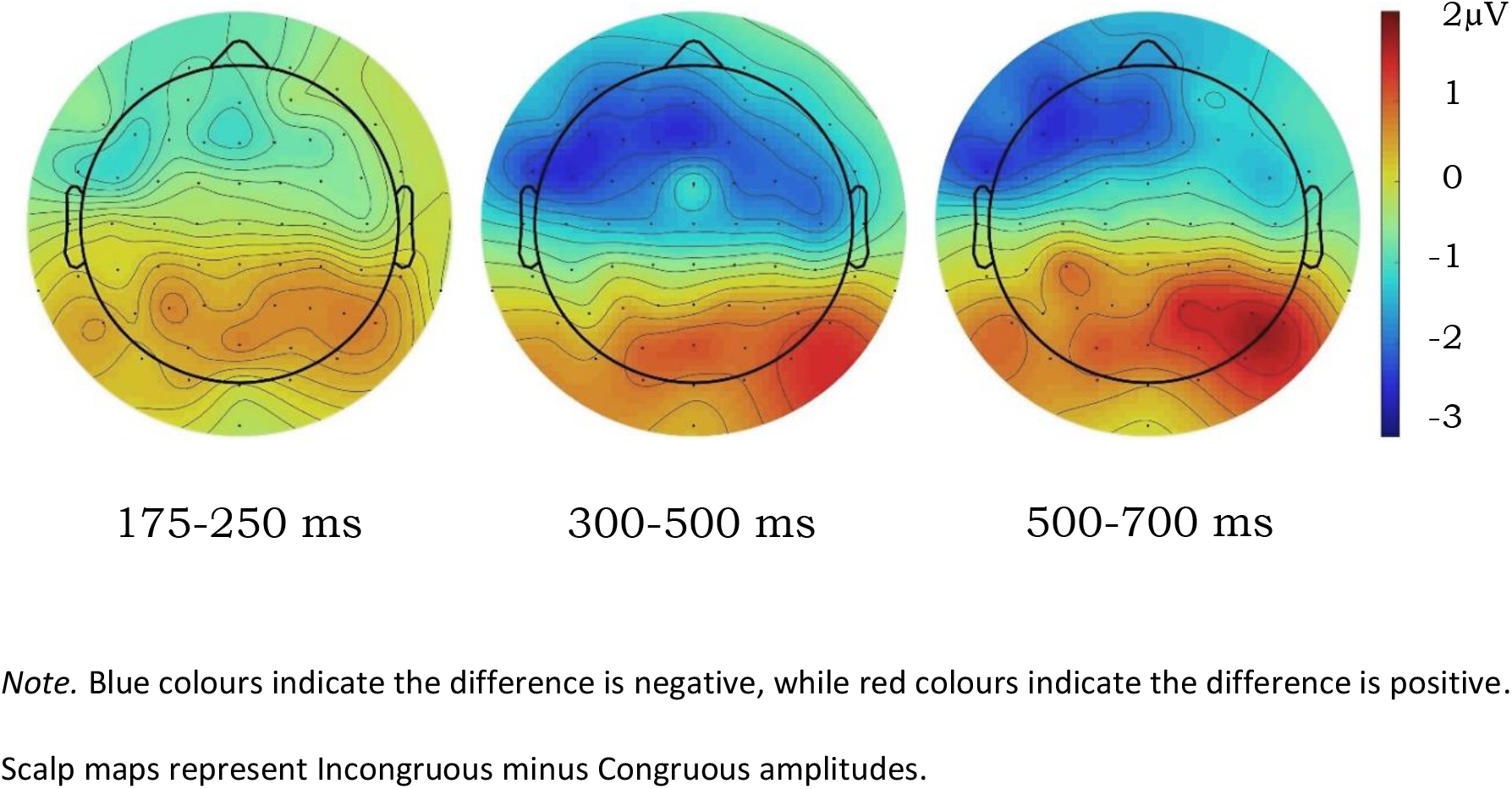
Scalp Maps of the Mean Voltage Difference Between the Congruency Conditions for Each of the Time Windows Under Investigation

**Figure 11.**
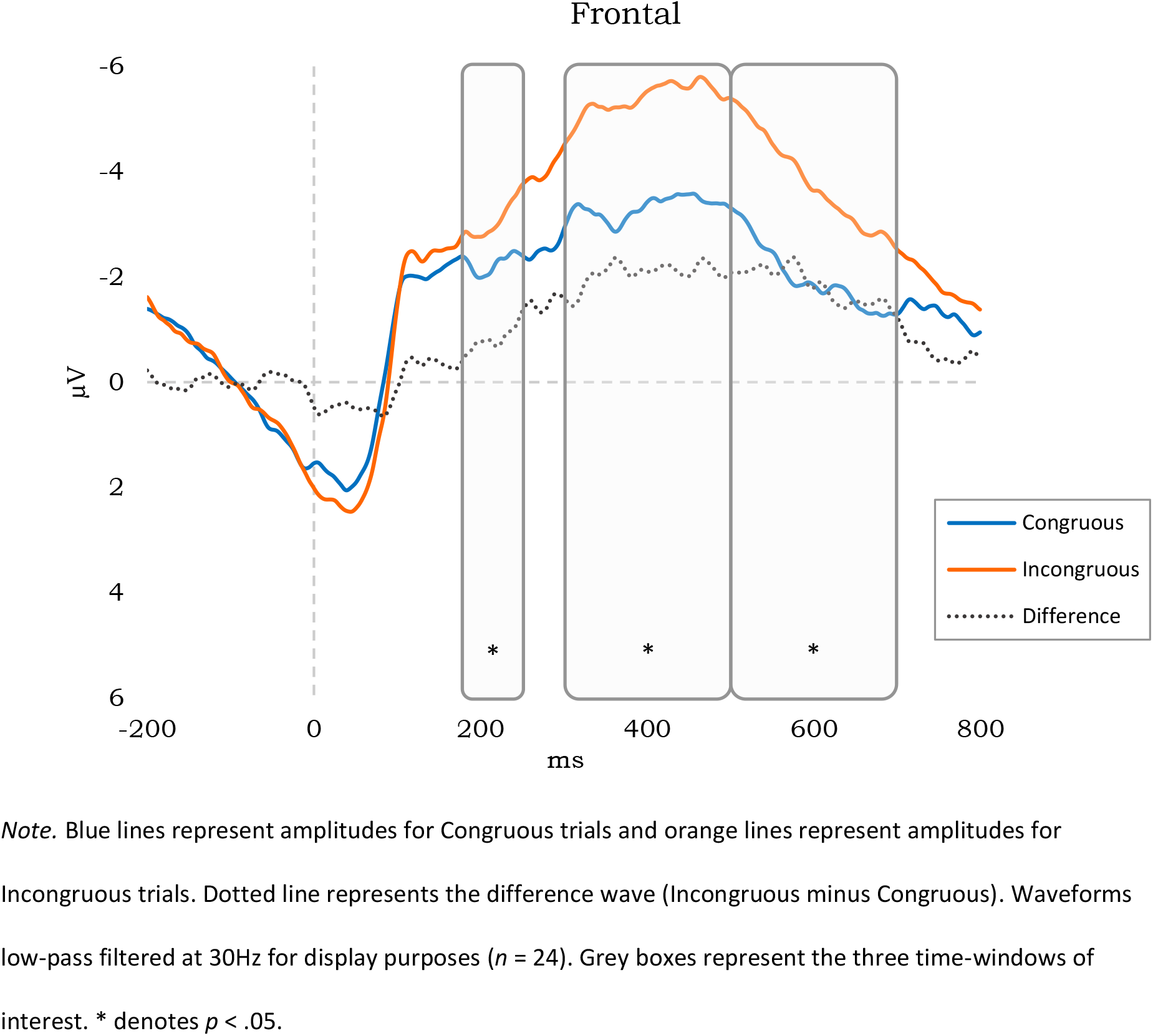
Grand-averaged ERPs for the Frontal Region, Collapsed Across Hemispheres

**Figure 12.**
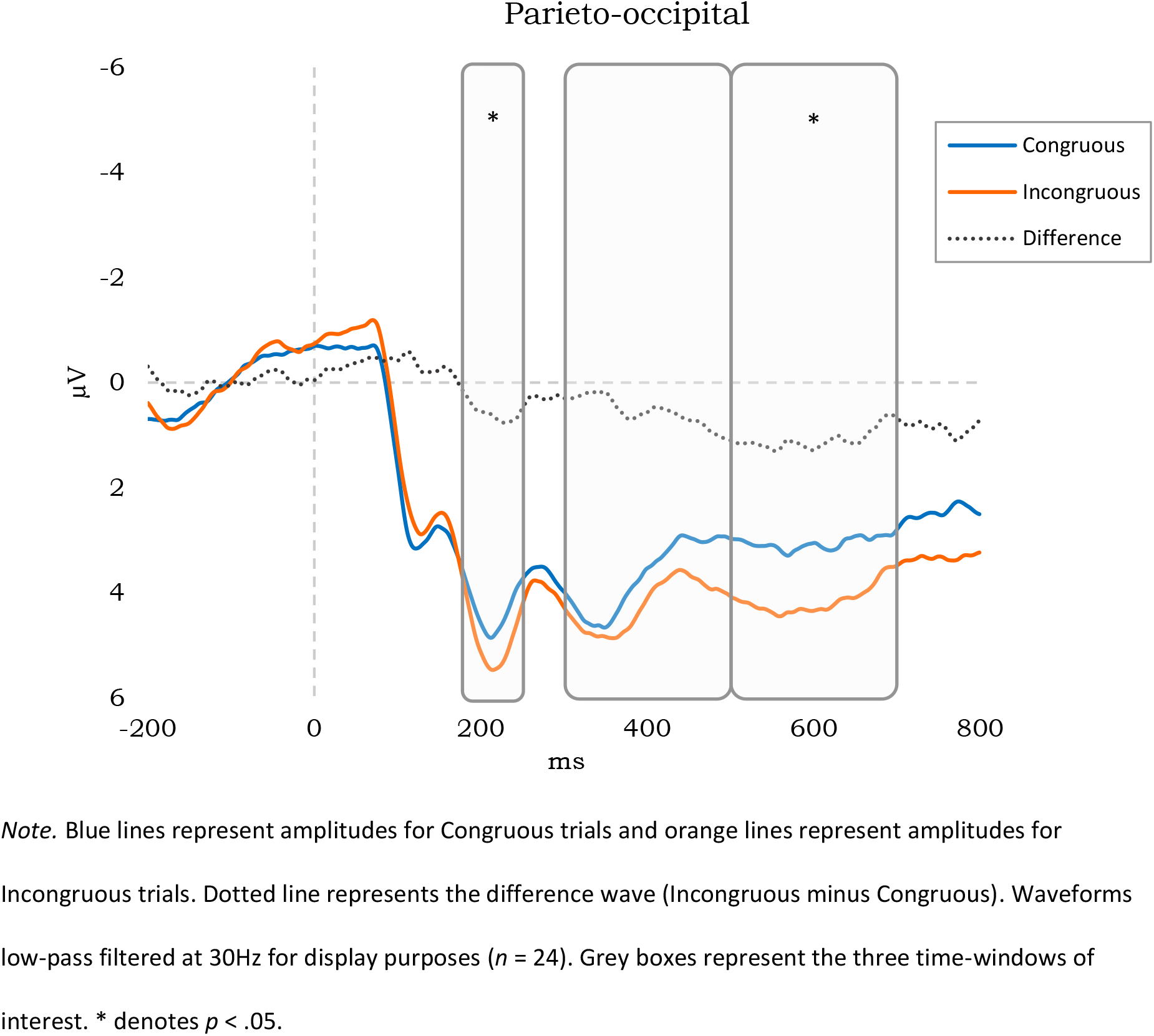
Grand-averaged ERPs for the Parieto-Occipital Region, Collapsed Across Hemispheres

### 300-500 ms Window

A three-way ANOVA revealed a main effect of Congruency, *F*(1, 23) = 6.16, *p* = .021, ƞp^2^ = .21, due to there being significantly more negative mean amplitudes for Incongruous trials (*M* = − 0.85) than Congruous trials (*M* = −0.02) during this time-window. There was no three-way interaction (*p* = .136), nor a Hemisphere x Region interaction (*p* = .274), nor a Hemisphere x Congruency interaction (*p* = .117). There was, however, a significant Region x Congruency interaction *F*(1.46, 33.65) = 32.92, *p* < .001, ƞp^2^ = .59. In terms of this interaction, follow-up paired t-tests revealed no significant effect of congruency within the Parieto-occipital region (*p* = .129). However, there was a significant effect within the Centro-parietal region, *t*(23) = 2.40, *p* = .025, *r* = .45. This represents a medium-to-large effect. This was due to there being a significantly more negative mean amplitude for Incongruous trials (*M* = −1.39 µV) than Congruous trials (*M* = −0.41 µV) within the Centro-parietal region during this time-window (see Figure 13 for the grand-averaged Centro-parietal ERPs). There was also a significant effect within the Frontal region, *t*(23) = 5.37, *p* < .001, *r* = .75, representing a large effect. This was due to there being a significantly more negative mean amplitude for Incongruous (*M* = −5.40) than Congruous trials (*M* = −3.33).

**Figure 13.**
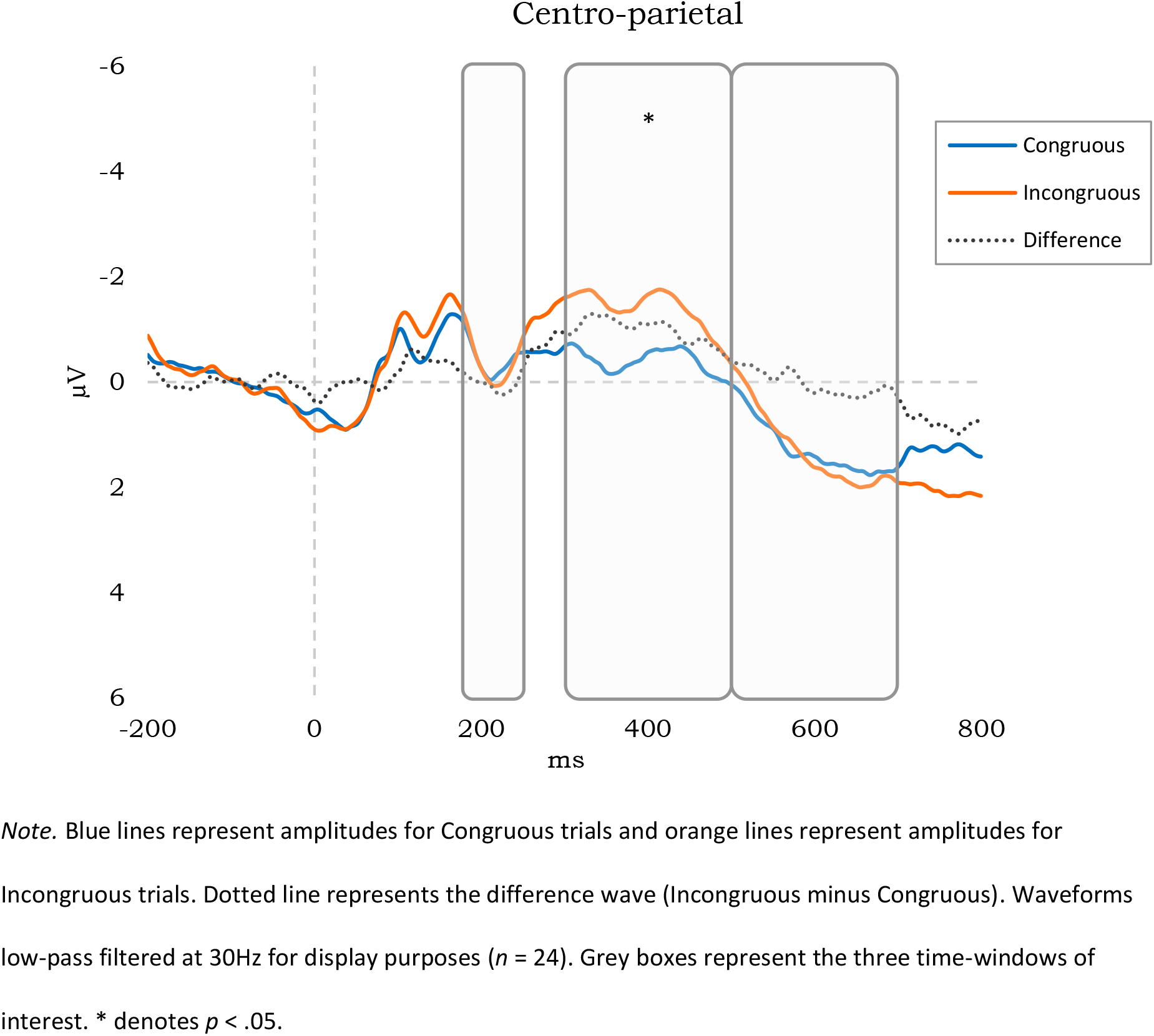
Grand-averaged ERPs for the Centro-Parietal Region, Collapsed Across Hemispheres

### 500-700 ms Window

A three-way ANOVA revealed no main effect of Congruency (*p* = .553). There was also no three-way interaction (*p* = .056), nor a Hemisphere x Region interaction (*p* = .656). There was, however, a significant Hemisphere x Congruency interaction, *F*(1, 23) = 5.72, *p* = .025, ƞp^2^ = .20.

Follow up paired t-tests for this interaction, collapsed across region, revealed no significant congruency-related difference in amplitude in either the left *(p* = .185) or right hemisphere (*p* = .958). There was also a significant Region x Congruency interaction *F*(2, 46) = 34.05, *p* < .001, ƞp^2^ = .60. In terms of this interaction, follow-up paired t-tests revealed no significant effect of congruency within the Centro-parietal region (*p* = .972), but did find a significant effect within the Parieto-occipital region, *t*(23) = −2.41, *p* = .025, *r* = .45. This represents a medium-to-large effect. This was due to there being a significantly more positive mean amplitude for Incongruous trials (*M* = 4.15 µV) than Congruous trials (*M* = 3.06 µV) within the Parieto-occipital region during this time-window. There was also a significant effect of congruency within the Frontal region, *t*(23) = 4.57, *p* < .001, *r* = .69, representing a large effect. This was due to there being significantly more negative mean amplitudes for Incongruous (*M* = −3.83) than Congruous trials (*M* = −1.99).

### ERP Results Summary and Discussion

Compared to Congruous trials, Incongruous trials displayed a significantly more positive mean amplitude for the P2 across the Parieto-occipital region, a significantly more negative mean amplitude for the N400 across the Centro-parietal and Frontal regions, and a significantly more positive mean amplitude for the P600 within the Parieto-occipital region. We also found significantly more negative amplitudes for Incongruous trials within the Frontal region across the early and late windows of interest.

The congruency-related amplitude changes seen in the P2 component help firm our contention that observer expectations were affecting gist processing, as this suggests top-down information was having an influence while perceptual processing was still ongoing. The sensitivity of the P2 to top-down information is still debated (e.g., Hansen et al., 2018), although previous work has relied on the presentation of individual images. It may be that changes to this early component seen here result from participants being able to generate expectations prior to target onset, thus providing the opportunity for more immediate top-down influence once the destination image is presented.

Amplitude changes across conditions were also apparent in the N400. There is a level of consensus that this component is a neural marker for semantic processing (see, for a review, Kutas & Federmeier, 2011), and so this finding supports our assertion that expectations were influencing higher-level cognitive processes rather than simply the extraction of low-level visual information. Furthermore, changes to the N400 have repeatedly been demonstrated for violations to semantic expectations within language (e.g., Van Petten, 1995) and across object-scene pairs (e.g., Demiral et al., 2012), and so the current findings suggest that an equivalent effect exists for violations between separate scenes. We found these N400 amplitude changes within the Centro-parietal as well as the Frontal region, in line with research showing the semantic processing of pictorial stimuli elicits a more anteriorly located negativity during this temporal window (e.g., Ganis et al., 1996). Further to this, we saw morphological dissimilarities within the ERPs across these two regions. In particular, the congruency-related amplitude differences in the anterior region spanned all three time-windows investigated, meaning the effect of approach-destination congruency was apparent in Frontal sites within 175-250 ms from target onset.

As with the N400, changes to the P600 again suggest an influence of higher-level processes on gist extraction. There is still debate, within both scene and language research, as to whether alterations to the P600 are a reflection of difficulties with semantic (e.g., Mudrik et al., 2010; Sitnikova et al., 2003) or syntactic (Hagoort & Brown, 2000; Võ & Wolfe, 2013) processing, or indeed an integration of both (Friederici & Weissenborn, 2007). Additionally, different aspects of syntactic processing have been proposed as being reflected in the P600 (e.g., Friederici et al., 2002; Gouvea et al., 2010; Kaan et al., 2000). For example, increased positivity at P600 has been elicited during ‘garden path’ sentences, which require a re-interpretation of expectations while reading a sentence due to an atypical grammatical format (Osterhout & Holcomb, 1992). Some equivalence to the current study is apparent, whereby expectations are built during a progression of approach images only to require re-evaluation once violated by the appearance of an incongruous destination.

In sum, differences were found in a putative scene-selective ERP component, related to integrating visual properties (P2), as well as later components related to contextual integration including semantic and syntactic coherence (N400 and P600, respectively).

## General Discussion

We conducted a series of behavioural and ERP experiments, involving the presentation of ‘approach’ images prior to target scenes. In so doing, we hoped to better understand the manner by which scenes are processed outside the laboratory, namely as elements of a progression of contextual information rather than simply isolated images. This allowed us to investigate whether semantic information was derived from the advancement through environments, and whether this generated ‘on-line’ expectations able to facilitate the processing of subsequent scenes. Experiments 1a and 1b investigated the effect of expectations on the processing of conceptual gist within scenes, through the manipulation of ‘approach-destination’ congruency, as well as the time-course of the effect through manipulation of target display duration. Experiment 2 then manipulated the sequentiality of these pre-target series, in order to investigate the influence of spatiotemporal coherence on gist processing, while Experiment 3 introduced a baseline condition to disentangle the separate roles of facilitation and interference on gist processing. Finally, in Experiment 4 we employed electroencephalography to chart the neural correlates associated with the manipulation of scene congruency.

As predicted, across experiments we found a benefit for categorising scenes when semantically congruous with lead-up images. Also in line with predictions, Experiment 1a revealed an advantage that was greatest at shorter target durations, where the opportunity to process visual information was most limited. This pattern of results was mirrored in Experiment 1b, where congruous target scenes only appeared on a quarter of trials, indicating the effect was not simply based on the frequency of conditions and that participants’ predictions as to an upcoming destination were being driven by an automatic mechanism. Next, Experiment 2 revealed that the performance advantage seen for Congruous trials was based on approach images providing a semantic context for upcoming targets. While a main effect of approach-image sequentiality was found, an increase in categorisation ability for Congruous-Sequential compared to Congruous-Disordered trials only neared significance, contrary to our predictions. Then, Experiment 3 confirmed that providing participants with semantically congruous approach images led to a facilitation of gist processing, as hypothesised, and demonstrated reduced performance compared to baseline when trials were incongruous in nature.

Finally, Experiment 4 investigated the neural correlates of predictability on rapid scene processing, showing an effect across all tested ERP components. For Incongruous trials, the P2 and P600 showed significantly greater mean amplitudes within the Parieto-occipital region, while a significantly more negative mean amplitude for the N400 was seen within the Centro-parietal and Frontal regions. Furthermore, Incongruous trials were also associated with a significantly more negative amplitude across the early and late time-windows within Frontal sites. Taken together, this meant we found congruency-related changes within the earliest known indicant of scene-specific processing (P2), within the component classically proposed as an index of semantic expectancy as well as the retrieval of conceptual information (N400), and within the component associated with both semantic and syntactic processing (P600). We will begin by addressing the findings from the behavioural experiments, before moving on to discuss potential interpretations for the task-related alterations to brain activity seen here.

Firstly, the condition-based differences in categorisation performance within Experiments 1a-b reveal that an observer’s expectations can alter scene processing. Importantly, the most substantial differences were found at target durations of 50 milliseconds and below, indicative of expectations influencing the earliest stages of processing. It appears, therefore, that top-down information has a role in modulating the extraction of scene gist. These results are perhaps not surprising when we consider the considerable quantity of research demonstrating an influence of expectations on the subsequent processing of the environment (Endsley & Garland, 2000; Langham et al., 2002; Mahon, 1981). Furthermore, such results appear to mirror the finding that the processing of objects is facilitated when contextually related to the scenes in which they are embedded (Antes et al., 1981; Biederman et al., 1973; Boyce & Pollatsek, 1992, Davenport & Potter, 2004; Underwood, 2005; Võ & Henderson, 2011). This has not only been found during the simultaneous presentation of scenes and their objects, but also when a scene is presented and then removed from view prior to the target object being displayed (Palmer, 1975).

So, just as the rapid processing of semantic information can influence the processing of objects, our results show the same is true for scenes. While ‘scene priming’ effects have been reported before (e.g. Sanocki & Epstein, 1997), such facilitation is likely based on low-level information, such as the priming of visual features (Brady et al., 2017) or the maintenance of basic scene layout in memory (Oliva & Torralba, 2001). The findings seen here, on the other hand, relate to the processing of scenes at the semantic level, as further discussed below, and so propose an expansion of the concept of scene priming. Just as a scene can be semantically primed by text displayed prior to its presentation (Reinitz et al., 1989), it appears that similar facilitation is possible when a target scene is preceded by a semantically relevant context. Such a finding adheres to the typical mechanisms underlying visual processing, where the constraints of cognitive capacity drive us to look for predictable patterns within the environment, from which to form expectations that lower the demands of subsequent processing (e.g., Bartlett, 1995; Gregory, 1997). In addition, it lies in agreement with recent findings that demonstrate pre-target narrative sequences are able to affect subsequent scene processing (Smith & Loschky, 2019).

The relationship between expectations and gist processing builds on complimentary work concerning improbable scenes (Greene et al., 2015). That research uncovered increased difficulty in understanding the meaning of atypical scenes, pointing to a disruption in gist processing when scenes diverge from what an observer expects. Such findings strongly point to a role of top-down information in rapid scene understanding, although there is an important distinction to our work. The violation of expectations within single scenes – such as a boulder inside a room (Greene et al., 2015) – would potentially result from inconsistencies between the bottom-up signal and a template stored in long-term memory. On the other hand, our study showed the effect of violating predictions based on the on-line flow of information: the introduction of approach images meant predictions could be formed prior to target onset, potentially resulting in the pre-activation of templates expected to be required for matching against the stimulus. Such pre-activation provides the opportunity for a stored representation to be available prior to the appearance of the target-derived signal, conceivably resulting in more rapid matching or, through predictive coding mechanisms, allowing for the detection of inconsistencies at an earlier processing level due to pre-emptive changes in error thresholds (Rauss et al., 2011).

These results stand in opposition to ‘forward sweep’ models, which assume minimal top-down modulation of gist processing (Itti et al., 1998; Potter et al., 2014; Rumelhart, 1970). However, we do not contend this necessarily rejects the primacy of bottom-up visual factors in scene perception. Across each of our behavioural experiments the accuracy with which scenes were categorised far exceeded chance level, even when approach images were incongruous with destinations. In other words, some degree of gist processing was still possible when no relevant semantic information was provided prior to destination-scene onset. Therefore, we propose that feature extraction mechanisms may well be capable of rapidly distinguishing a great deal of information within complex natural scenes (e.g., Potter et al., 2014), but that these mechanisms are susceptible to influence from higher-level processing. This might particularly be the case when, as in our design, antecedent information is provided before gist processing begins, thereby allowing for the formation of expectations prior to a scene being encountered. So, we cannot comment on the processing of individual, segregated scenes, as we did not investigate this. What we do contend, however, is that under conditions which better reflect functioning outside the laboratory it appears that top-down information, in the form of expectations, affects conceptual gist processing.

Secondly, the results from Experiment 2 show the performance advantage for Congruous trials was largely due to the provision of contextual information. More specifically, the facilitation of gist processing through expectations appears to be driven by the observer being provided with semantic information, in the form of an environmental setting, from which more accurate predictions can be formed. Perhaps surprisingly, the sequentiality of the approach images did not significantly influence performance, and so we found no evidence that the spatiotemporal coherence of series, or the generation of a perceived flow of movement, had a bearing on participants’ categorisation ability. This was counter to our predictions, based on recent work identifying an important role for the narrative coherence of pre-target sequences (Cohn et al., 2012; Smith & Loschky, 2019).

While we did find evidence for a significant congruency-sequentiality interaction, as well as a significant main effect of sequentiality, the expected effect within Congruous trials only neared significance. This suggests that – within the particular constraints imposed by our design – if an effect of sequentiality does exist its influence is much reduced as compared to the effect of congruency. However, this is not necessarily true under all circumstances, and there is an important distinction from previous work. The approach images adopted here were within relatively close proximity to their eventual destination, and so the narrative created by these series is not comparable to the narrative created across the panels of a comic strip (Cohn et al., 2012), nor the strings of images depicting a journey from one distinct location to another that is spatially distant (Smith & Loschky, 2019). So, the generated story of “I am approaching a shop” may have remained unaltered irrespective of whether the leading images were sequential or not, something that would be unlikely for the more complex narratives within previous designs. However, while the narrative of our sequences may not have been overly disrupted by being disordered, this manipulation certainly disrupted the appearance of linear movement. As a consequence, our results find there to be comparatively minimal additional benefit from the creation of ‘perceived flow’ (e.g., Gibson, 1966).

A final point to make relating to the results of Experiment 2 is that they help alleviate any concern regarding the origin of the congruency-based changes in performance. Our design meant some target scenes necessarily contained low-level similarities with the most proximal leading images. For instance, on a congruous (and sequential) trial the final leading image of an approach to a shop may allow for some information relating to the final scene to be pre-empted because, say, the general dimensions of the store could be determined by a partial view through the window. For this reason, it might be argued that the categorisation performance changes across congruency conditions were due to the priming of low-level visual features prior to target onset, whether these would be similarities in terms of constituent features (Shafer-Skelton & Brady, 2019), layout (Sanocki & Epstein, 1997) or ‘spatial envelope’ (Oliva & Torralba, 2001). However, in Experiment 2 the disordered nature of approach images meant targets were immediately preceded by an image more spatially distant to the destination, with the implication that the similarities in low-level information across the final leading image and the target scene were, by definition, reduced. Despite this, the effect of congruency remained.

While the preceding behavioural experiments provided clear support for an effect of expectations on gist processing, it was important to investigate the manner of such influence. As the previous iterations did not contain a control condition it remained open to question whether gist processing was being facilitated by congruous approaches, inhibited by incongruous approaches, or a mixture of both. Therefore, in Experiment 3 we introduced a ‘No-context’ condition where approach images were replaced with images of coloured patterns, allowing us to maintain the same trial structure while removing any pre-target semantic information. By doing so, this condition served as a measure of baseline performance, in terms of gist processing ability in the absence of antecedent contextual information, to which the congruency conditions could be compared. As predicted, we saw significantly increased categorisation ability on Congruous trials when compared to baseline performance, confirming that contextual information facilitated subsequent gist processing. Such a finding was expected due to the well-understood mechanisms of visual processing, where increased efficiency is achieved through utilisation of learned regularities to generate expectations as to the current environment (Chaumon et al., 2008; Fiser et al., 2016; Gregory, 1997; Li et al., 2004; Roc, 1997; Ullman, 1980).

While we did not make predictions as to whether a cost to processing on Incongruous trials would be apparent, Experiment 3 also demonstrated interference to gist extraction when participants were provided with inappropriate contextual information. This appears to be in agreement with previous gist processing research, where improbable scenes – i.e. those which contain unexpected features – were found by participants to be more difficult to extract meaning from, as compared to typical scenes (Greene et al., 2015). It is similarly in line with a recent investigation using object-scene pairs, which showed not only contextual facilitation of object processing but also interference to performance as a result of semantic violations within pairs (Lauer et al., 2020).

The exact mechanisms governing such interference remain open to interpretation, although it seems reasonable to suggest that the deficit in performance results from an attempt to match an unexpected bottom-up signal to an inappropriate, internally generated representation. This may be in the form of predictive coding mechanisms, whereby a significant disparity between expectations and ascending signal leads to prediction errors substantial enough to force reanalysis of the sensory input (e.g., Barrett & Simmons, 2015; Macpherson, 2017; Talsma, 2015). Alternatively, the Scene Perception and Event Comprehension Theory (SPECT; e.g., Loschky et al., 2018) proposes that an observer creates an internal current event model while progressing through a narrative, which represents their understanding of what is happening in that moment. Within this framework, significant changes in situational continuity initiate an automatic cognitive shifting towards creation of a new event model, and this operation is associated with distinct processing costs (Loschky et al., 2018). In the current study, therefore, reduced performance may have resulted from the disruption to processing due to the break in contextual continuity within Incongruous series. On the other hand, the case could be made that participants continued to search for an associative link when confronted with the lack of coherence within Incongruous trials, resulting in a protracted cognitive load that affected low-level perceptual processes (Afiki & Bar, 2020), or even that violations to predictions invoked increased encoding of the current scene-image while actively suppressing retrieval mechanisms (Sherman & Turk-Browne, 2020).

Turning to Experiment 4, changes to the neural signature help elucidate both the time-course and means by which the violation of expectations affects processing. Firstly, the contention that observer expectations were influencing gist processing is further strengthened by the display of changes to early ERPs, specifically the congruency-related amplitude differences in the P2 component. Changes in amplitude appearing so soon after target onset suggest an influence of top-down information while perceptual processing was still ongoing, similar to that proposed for object recognition (Bar, 2003; Fenske et al., 2006). While the P2 has previously been advanced as a marker for scene processing (Harel et al., 2016), there is debate as to whether this component is sensitive to top-down influence. For example, recent research found no top-down modulatory effect (Hansen et al., 2018), at least in relation to observer-based goals. Conversely, some forms of early higher-order influence have been implied, as changes to amplitude at ∼200 ms post-stimulus have been observed with tasks involving the detection of objects within natural scenes, potentially reflecting decision-related activation (Thorpe et al., 1996; VanRullen & Thorpe, 2001), and tasks that manipulated the emotional nature of scene-images, argued as being driven by motivational systems (Schupp et al., 2006).

It is possible that different forms of top-down information are integrated at different temporal points, or simply that modulations to such early ERP components are more apparent under certain experimental designs than others. It may be that changes to the P2, found here, result from the use of antecedent information. Our use of approach images allowed for an expectation of the upcoming target category to be formed prior to its onset, meaning that this top-down information was available to facilitate processing from the moment the destination scene was presented. This is a clear departure from a task that involves a single image, whereby bottom-up input, perhaps in terms of low spatial frequency information (e.g., Bar et al. 2006), has to first be employed at scene onset to form expectations and only then is available as a tool for the ongoing evaluation of the incoming signal. As a result, it appears reasonable that a design eliciting expectations prior to target-onset would be able to more swiftly affect early ERP components such as the P2, as compared to single-image designs. Moreover, this likely better reflects processing during day-to-day life, where we constantly generate expectations as to the setting we are to encounter next (e.g., Bartlett, 1995).

Secondly, condition-related changes in the magnitude of the N400 component suggest the processing of Incongruous trials was affected by perceived semantic violations within those series. Evidence has repeatedly indicated that the N400 is a neural marker for semantic processing, with increased negativity in this temporal window being related to difficulties with semantic integration (Barrett & Rugg, 1990; Kutas & Hillyard, 1980; McPherson & Holcomb, 1999). Our display of increased negativity at the N400 mirrors previous work related to the semantic violation of object-scene pairs. Such effects have been observed both when a scene is presented prior to target-object presentation, thereby allowing for *a priori* expectations as to the identity of the upcoming object to be formed (e.g., Demiral et al., 2012; Ganis & Kutas, 2003; Võ & Wolfe, 2013), as well as during simultaneous presentation of objects and scenes, meaning expectations as to object appropriateness cannot be formed prior to onset (e.g., Mudrik at al., 2010). However, while this previous work investigated violations to the semantic relationship between single scenes and their objects, our results show a comparable neural signature resulting from semantic violations between scenes. Additionally, that the experimental manipulation led to alterations in the N400 again makes it improbable that effects were due to confounds based on the repetition of low-level features. This component reflects a later stage of processing (e.g., S. Wang et al., 2017), its association with semantic integration, across multiple modalities, is much replicated (for a review, see Kutas & Federmeier, 2011), and its origins have been localised to higher-order brain regions such as those involved in semantic unification processes (e.g., Lau et al., 2008; L. Wang et al., 2012).

The processing of semantic information related to images, as opposed to text, has often been shown to elicit a more anterior negativity during this temporal window (e.g., Ganis et al., 1996; Holcomb & McPherson, 1994; Kutas et al., 2006), and our results reflect this. However, while both Frontal and Centro-parietal sites here displayed typical N400 effects, in terms of increased negativity for Incongruous trials, the pattern of amplitude changes are morphologically dissimilar across regions. Notably, the congruency-based amplitude changes in anterior sites began to emerge earlier (∼200 ms) and were sustained for a far greater period of time (until at least 750 ms after target onset), with significantly more negative amplitudes for Incongruous trials across all three time-windows. There is minimal research regarding similar late effects at anterior sites, although it has previously been attributed to late processes of semantic evaluation (Mudrik et al., 2014).

On the other hand, investigations of pre-N400 negativity across frontal regions have been more frequent. In particular, the earlier emergence of effects at anterior compared to central sites has repeatedly been observed in object-scene research, leading to the proposition that this reflects a separate component, namely the N300 (Barrett & Rugg, 1990; Demiral et al., 2012; McPherson & Holcomb, 1999; Truman & Mudrik, 2018). This has been offered as reflecting context effects at a perceptual level (e.g., Schendan & Kutas, 2002; Mudrik et al., 2010), immediately prior to the semantic processing indicated by the subsequent N400. Furthermore, the N300 appears to be sensitive to alterations in global stimulus features rather than to low-level visual elements (e.g., Schendan & Kutas, 2007), and recent work has suggested it may be an index of perceptual hypothesis testing at a scale of whole scenes and objects, such as template matching routines based on perceptual structure (Kumar et al., 2020). It has also been put forward that components prior to the N300 may reflect predictive coding mechanisms in relation to expected low-level visual features (Kumar et al., 2020). However, distinguishable N300 effects have often not been forthcoming (e.g., Demiral et al., 2012; Ganis & Kutas, 2003) and this dissociation between the N300 and N400 is still debated (see, for example, Draschkow et al., 2018; Willems et al., 2008).

It is important to note that our early window of interest (175-250 ms) preceded the window typically used for investigating the N300 (e.g., Kumar et al., 2020; Lauer et al., 2020), and so our intention is not to comment directly on the debate surrounding that particular component. What we do assert, however, is that – if the N300 is taken as indexing perceptual, rather than higher-order, processing – then our early effects across anterior regions should be similarly categorised. In other words, due the early amplitude changes within Frontal sites as well as the alterations to the P2 discussed above, we suggest that expectations generated prior to target presentation were able to influence the extraction of scene gist at the level of perceptual processing. Predictions as to the category of an upcoming scene are likely to contain predictions not just of its identity, but also its expected perceptual features. At one level an observer may expect to see a beach, but on another level they may be expecting a certain spatial layout (Sanocki & Epstein, 1997) or specific form of spatial envelope (Oliva & Torralba, 2001), or a certain array of colours (Castelhano & Henderson, 2008; Gegenfurtner & Rieger, 2000), textures (Renninger & Malik, 2004), edge-based information (Walther & Shen, 2014) or other low-level features (Shafer-Skelton & Brady, 2019). However, whether the expectation-based violations to processing seen here were related to global properties or to lower-level information remains open to debate.

In terms of the P600, the changes observed here help further elucidate the potential mechanisms underlying the effect of expectations on scene processing. As with the N400, previous scene-related studies investigating this component have focused on object-scene pairs, but findings have proved inconsistent. For instance, Mudrik and colleagues (2010) found that positioning inappropriate objects in appropriate places (a semantic violation, such as a chessboard – rather than a baking tray – being placed into an oven) led to a more negative amplitude at 600 ms, compared to scenes containing appropriate objects. Võ and Wolfe (2013), alternatively, found no alterations to the P600 with similar object-scene semantic violations, but did find an increased P600 when appropriate objects were presented in a position considered to be atypical (such as a dishtowel on the floor, as opposed to hanging on a nearby towel rail). The authors proposed that these images created syntactic – rather than semantic – violations, as the objects contravened structural rules while remaining semantically congruous with their scenes. Thus, they reported the P600 as reflecting syntactic violations to scene processing (Võ & Wolfe, 2013), and so there appears to be a lack of consensus regarding the types of context-based violation that lead to changes in this component. However, it may be the case that these differing results reflect sensitivity to different methodological choices across studies, such as whether the scene is presented prior to the object or simultaneously with it, and whether the object is in a position of stable rest or being acted upon by agents within the image.

The current study, on the other hand, found alterations to the P600 without such violations to object location or appropriateness. It may be the case, therefore, that these similar ERP patterns are reflecting different phenomena, as research has shown the P600 to be associated with different forms of syntactic anomaly (Gouvea et al., 2010). Increased positivity at the P600 for inconsistent syntax between scenes and objects may be akin to grammatical errors in sentences (e.g., Hagoort et al., 1993), whereas the increased positivity seen here might be more similar to that elicited by ‘garden path’ sentences (e.g., Osterhout & Holcomb, 1992). Although containing no grammatical errors, progression through such sentences reaches a point where re-interpretation of expectations is necessary, through parsing the word-sequence in a different way. A similar form of violation may be responsible for our P600 pattern, whereby the progression of sequential approach images built an expectation in the observer – much like the expectation created during progression through the words of a sentence – until the final, incongruous destination disrupted the assumed end-point and resulted in an attempted re-evaluation of meaning. So, it may not be the case that the P600 is exclusively within the purview of violations to syntax, as it could also be a marker of the sudden need for reanalysis elicited by the disruption to an expected sequence. Such an explanation remains speculative, and further work surrounding the similarities in neural signatures across scene processing and language comprehension is certainly warranted. Both the N400 and P600 in scene processing appear somewhat analogous to those from language comprehensions studies and, while the specific forms of ‘grammar’ involved in these differing tasks likely diverge, a strong case can be made for the existence of commonalities (e.g., Võ et al., 2019).

This is an inchoate area of research and alternative interpretations as to the mechanisms responsible for such effects are possible. What seems a reasonable proposition, however, is that antecedent information allowed for expectations as to the category of the upcoming scene to be automatically generated. These expectations could be used to pre-activate internal representations or templates of expected-category exemplars which then become available for matching against the target scene once presented. Certainly, there are many separate conceptualisations of perceptual hypothesis testing which could be applied to our findings (see, for a review, Clark, 2013), although where such matching might take place within the visual processing stream remains open to debate. As we found congruency-based alterations to the ERP across all time-windows of interest, this recommends that it may be unwise to envisage a singular temporal or cortical point at which top-down predictions affect processing.

On one hand, our finding of early expectation-based amplitude changes indicates that predictions did influence feature extraction mechanisms. This is in line with recent findings pointing to a role of top-down feedback in the earliest stages of perceptual processing. Research using fMRI and multivariate pattern analysis has shown that expectations as to an upcoming, non-complex visual image are able to evoke stimulus templates in the primary visual cortex (Kok et al., 2012; Kok et al., 2014), and specifically within those deep layers proposed as being responsible for sending feedback to upstream regions (Aitken et al., 2020). Relatedly, higher-level cognitive factors have been shown to affect neurons in early sensory cortex (Lamme & Roelfsema, 2000), while representations based on semantic content have been shown to influence the extraction of elementary image features (Neri, 2014). It should be noted, though, that the semantic control of early sensory processing is still debated (see, for example, Carandini et al., 2005; Heeger et al., 1996).

On the other hand, our pattern of results reveals the violation of expectations also had an effect at a more advanced level of the processing stream. This contention is based on a number of factors. Firstly, the pattern of neural responses relating to the later components closely mirrors those long-associated with higher-order processing (e.g., Barrett & Rugg, 1990; Demiral et al., 2012; Ganis & Kutas, 2003; Kuperberg, 2007; Kutas & Hillyard, 1980; Lau et al., 2008; McPherson & Holcomb, 1999; L. Wang et al., 2012). Secondly, previous work showing the importance of expectations on gist processing found considerable deficits in the processing of improbable real-world scenes even when matched to probable scenes in terms of their low-level visual features, strongly implying that expectations were affecting processing at a stage somewhat beyond the level of initial feature extraction (Greene et al., 2015). Lastly, the superordinate category of targets in the current study (in terms of interior / exterior distinction) was maintained during Incongruous trials, ensuring that there was similarity across the low-level information present. For instance, a retail store scene may contain much of the same general structure or non-localised amplitude information as that of a supermarket, in terms of openness, roughness, etc. However, it should be noted that the exact level of similarities in low-level information across both superordinate and basic level scene categories is debated (e.g., Banno & Saiki, 2015; Fei-Fei et al., 2007; Gerhard et al., 2013; Loschky & Larson, 2008; Oliva & Torralba, 2001).

Taken together, it appears that *a priori* expectations had a broad effect across multiple stages of scene processing. Indeed, the concept of having a specific point of effect is perhaps only valid if a linear hierarchy of visual processing is accepted, as opposed to a cognitive network displaying abundant re-entrant connections (e.g., Boehler et al., 2008; Bullier, 2001; Koivisto et al., 2011). It may be, therefore, more germane to think of predictions of an upcoming scene as influencing manifold areas within the hierarchy simultaneously, whereby expectations set a cortical ‘state’ deemed appropriate for processing the predicted upcoming signal across the whole network (Gilbert & Li, 2013). Such an account could be considered as fitting within predictive coding frameworks (e.g., Friston, 2010; Friston & Kiebel, 2009; Rao & Ballard, 1999). Internal representations, activated through expectations as to the upcoming scene category, could allow for top-down predictions to propagate across processing areas (e.g., Lewis & Bastiaansen, 2015). As such, regions are informed by predictions based on the approach images, where reanalysis becomes necessary if the bottom-up signal is fundamentally at odds with what was expected (e.g., Talsma, 2015). In other words, a significant discord between predictions and input may create a substantial prediction error that crosses a pre-determined threshold or criterion, forcing both a major update of the internal model and reprocessing of the sensory signal (e.g., Barrett & Simmons, 2015; Macpherson, 2017; Talsma, 2015). Importantly, under such a model, *a priori* expectations may alter prediction error thresholds not only in early visual areas but also within higher-order processing regions (e.g., Hindy et al., 2016; Huang & Rao, 2011; Lewis & Bastiaansen, 2015; Summerfield et al., 2006), thus potentially resulting in a situation where difficulties in matching become apparent across separate levels of abstraction, such as at a perceptual and conceptual level.

The current study opens several important lines for further investigation. The finding that low-level visual information apparent at stimulus onset is not the only influence on gist processing asks the question as to what other sources of influence might exist. These could range from differing forms of top-down communication, such as an observer’s goals, to the role of other sensory information, such as potential cross-modal facilitation through the parallel presentation of visual scenes and their related sounds. The design employed here attempted to better reflect scene processing outside the lab, but there are limits to how immersive a series of static images can be. To take this a stage further, leading images could be replaced by video clips of journeys or, better still, the incorporation of VR technology could embed participants within pre-determined environments. An important question that remains concerns the precise nature of the spatiotemporal dynamics of expectation effects. Other methodologies might be able to offer insights, such as the use of transcranial magnetic stimulation for interrupting re-entrant communication, or the application of dynamic causal modelling to tease apart the respective roles of top-down and bottom-up information. Finally, there appears to be clear similarities with how expectations affect the processing of meaning across both scenes and language. However, more work is needed to uncover the true extent of these commonalities, such as whether this demonstrates a single, amodal semantic system in operation.

This study moved away from the traditional RSVP approach – and towards more ecologically valid scenarios – through the incorporation of ‘approach’ images prior to target-scene onset, and the findings presented here reveal an important role of expectations during scene processing. Specifically, predictions as to an upcoming scene, generated automatically, were able to facilitate processing when valid, and interfere with processing when invalid. Furthermore, the use of both behavioural and neuroimaging methods adds to our understanding of the temporal dynamics of rapid scene processing and indicates an influence of top-down communication on the extraction of conceptual gist. This runs contrary to models supposing exclusive analysis of low-level information as determining the processing of scene-gist, such as ‘forward sweep’ frameworks. In addition, we also put forward a case that *a priori* expectations are able to affect gist processing at both a perceptual and conceptual level. While the precise mechanisms by which expectations affect the processing of scenes are still to be discovered, we argue that semantically relevant antecedent information may allow for scene-category templates to be pre-activated across various areas within the visual hierarchy. Future insights may be forthcoming from research concerning predictive coding, which offers a potential framework for the utilisation of top-down information within the brief timeframes where gist processing takes place.

## Appendix

## Appendix A

### Creation of Image Series

The intention when constructing the series was to create progressions which mimicked movement through an environment towards a destination, while reducing instances of over-similarity across viewpoints and avoiding sudden ‘jumps’ in the progression. Accordingly, variations in geographical distances between approach images needed to be considered across series, mainly due to the differing constraints imposed by the superordinate categories. For example, the distance between points during a progression through a house to, say, a bedroom would be inherently shorter when compared to the points of progression towards a beach. An approach to a bedroom might begin with a view of a stairway across an atrium, followed by an image on the stairway, one at the top of the stairway turning onto a hallway, another at the mid-point of a hallway, and one turning the corner to show a bedroom doorway prior to the target being shown. In doing so, each transition of the approach would be accounted for, although the geographical distance covered would be relatively short. If the progression was towards a beach, on the other hand, then mirroring the distances between approach images from the bedroom series would result in five very similar viewpoints, almost indistinguishable from one another under the processing constraints of rapid presentation. As a consequence, in such instances we somewhat ‘stretched out’ the approach, so that it covered a greater geographical distance but at the same time maintained the principle of showing each transition in the journey, say from a carpark, down a pathway and between dunes before arriving at the beach. Again, care was taken to avoid sudden jumps in the narrative, so that the spatiotemporal relationship between successive leading images always remained apparent. This could be considered an attempt to instil a ‘semantic flow’ within each series, with each of the transitional points of the approach represented in a manner which maintained the sense of a progression throughout.

There are several other important points to note relating to the construction of series. Firstly, the destination scene could not be immediately determined from the earliest leading images. This was due to there being similar progressions across many series, in both interior-destination series (for instance, ‘bathroom’ and ‘bedroom’ targets would have similar approaches, involving stairways, hallways, etc.), and exterior-destination series (where many progressions shared similarities, such as traversing pavements, pathways and carparks). Furthermore, the eventual superordinate category of the target could not be anticipated at the start of the series: the approach images might represent a journey out in the open but with an indoor destination scene, or vice versa, such as walking across a garden before entering an outbuilding. Additionally, approaches frequently passed through other target categories. For example, images of a high street – a target category on some trials – might be passed through within the approach images of a series with a ‘shop’ target. It should be reiterated that this potential interplay across trials was at the category level, not the exemplar level, as no scenery (whether approach image or destination) was repeated at any point during the task.

Secondly, a balance had to be struck in terms of the final approach image representing a viewpoint geographically close enough to heighten expectations as to the destination, while trying to minimise the amount of similarity in low-level features across these two images. This was to ensure that performance was based on semantic prediction rather than simply on the repetition of low-level visual information. Therefore, while some features of a destination might be visible within the later approach images (such as the ocean on the horizon while progressing towards a ‘beach’ target, or the corner of a table and chair seen through a doorway prior to reaching a ‘dining room’ target) care was taken to maintain substantial differences in both the viewpoint and available visual features between the approach images and the destination scene. This practice was considered in line with the overarching tenet driving the construction of each series, namely that the progressions should mirror as closely as possible how individuals experience the environments in which they are embedded through the course of daily life.

Thirdly, the inclusion of people within images was kept to a minimum. It was not considered necessary to exclude pedestrians, shoppers, etc. from the sequences, as the aim was to represent environments in their usual state. However, care was taken to ensure that individuals within sceneries did not become a distraction from the experimental task, and so no images included people positioned close in the foreground or looking directly at the observer. Finally, all images (with the exception of multi-storey carparks) were of sceneries outside the county of the university’s location, in an attempt to limit any potential confounds due to familiarity with the specific exemplars used.

## Appendix B

### List of Scene Categories

ART GALLERY; BATHROOM; BEACH; BEDROOM; CARPARK; CHURCH; DINING ROOM; ENTRANCE HALL; FIELD; GARDEN; GRAVEYARD; HIGH STREET; KITCHEN; LIVING ROOM; MULTISTOREY CARPARK; OUTBUILDING; PARK; PETROL STATION; PUB; QUAY; RECYCLING AREA; RETAIL STORE; RIVER; ROAD; SHOP; SPORTS PITCH; SUPERMARKET; TAKEAWAY; TRAIN STATION; WOODS

## Appendix C

### Statistical Analyses

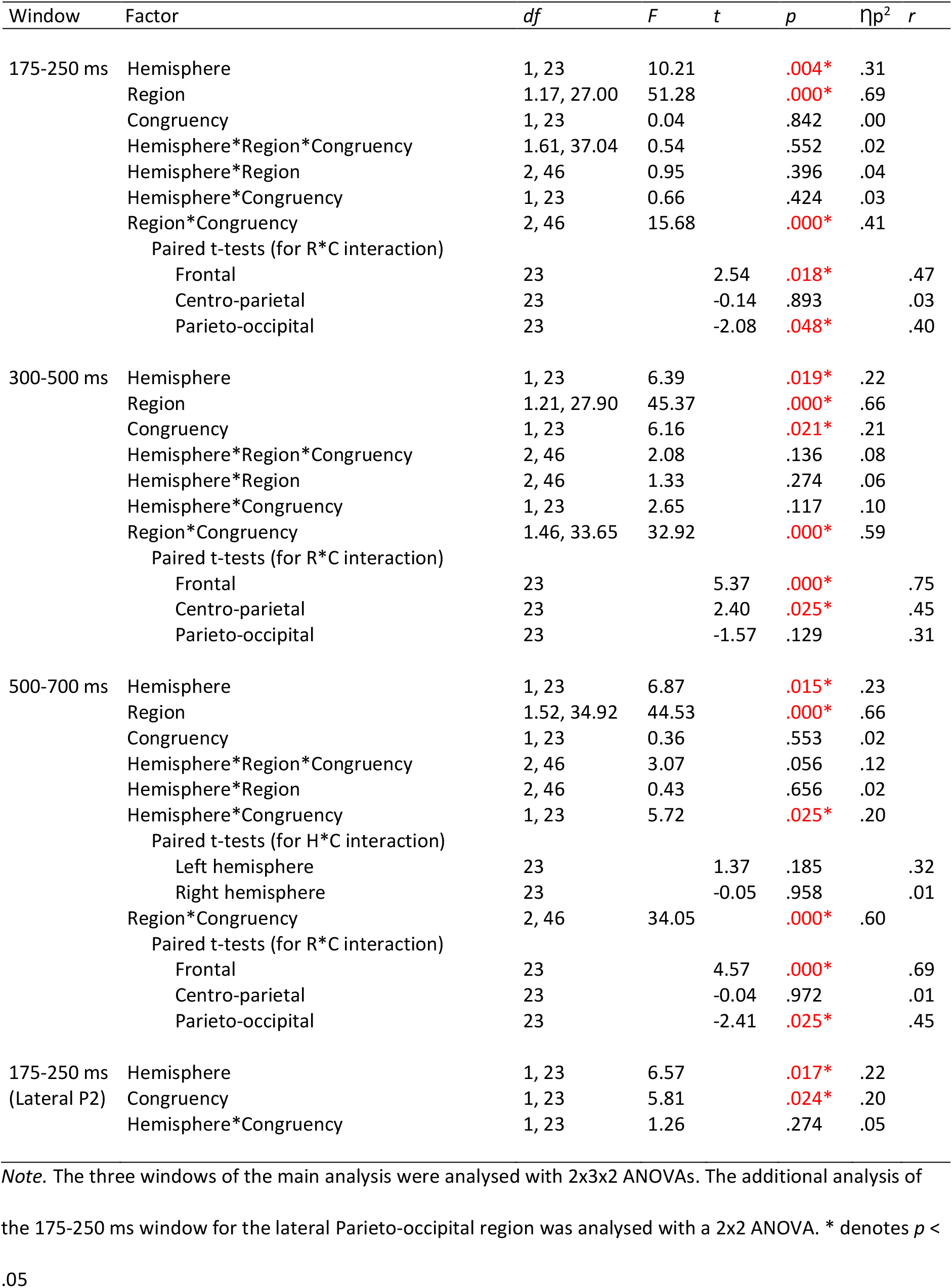

## Appendix D

### Additional Analysis: Lateral P2

In our initial analyses we found a significant effect of Congruency within the P2 time-window at posterior sites. However, in the interest of completeness we decided further investigation would be insightful. Previous scene processing research concerned with the P2 component has found effects to be maximal at sites more lateral than our initial ROIs (Hansen et al., 2018; Harel et al., 2016; Harel et al., 2020). Consequently, we created a Lateral Parieto-occipital ROI comprising six electrodes (split equally across hemispheres). The position of these regions was chosen to mirror previous work as closely as possible. Specifically, Harel and colleagues (2016; 2020) use a lateral region including eight electrodes across the two hemispheres (P5/P6, P7/P8, P9/P10 and PO7/PO8). Exact duplication of this setup was not possible, as instead of the electrode pair P9/P10 our array included TP9/TP10, which were located near the mastoids, and had been used as our re-referencing electrodes. Therefore, our lateral regions consisted of P5/P6, P7/P8 and PO7/PO8 (see Figure D1).

**Figure D1.**
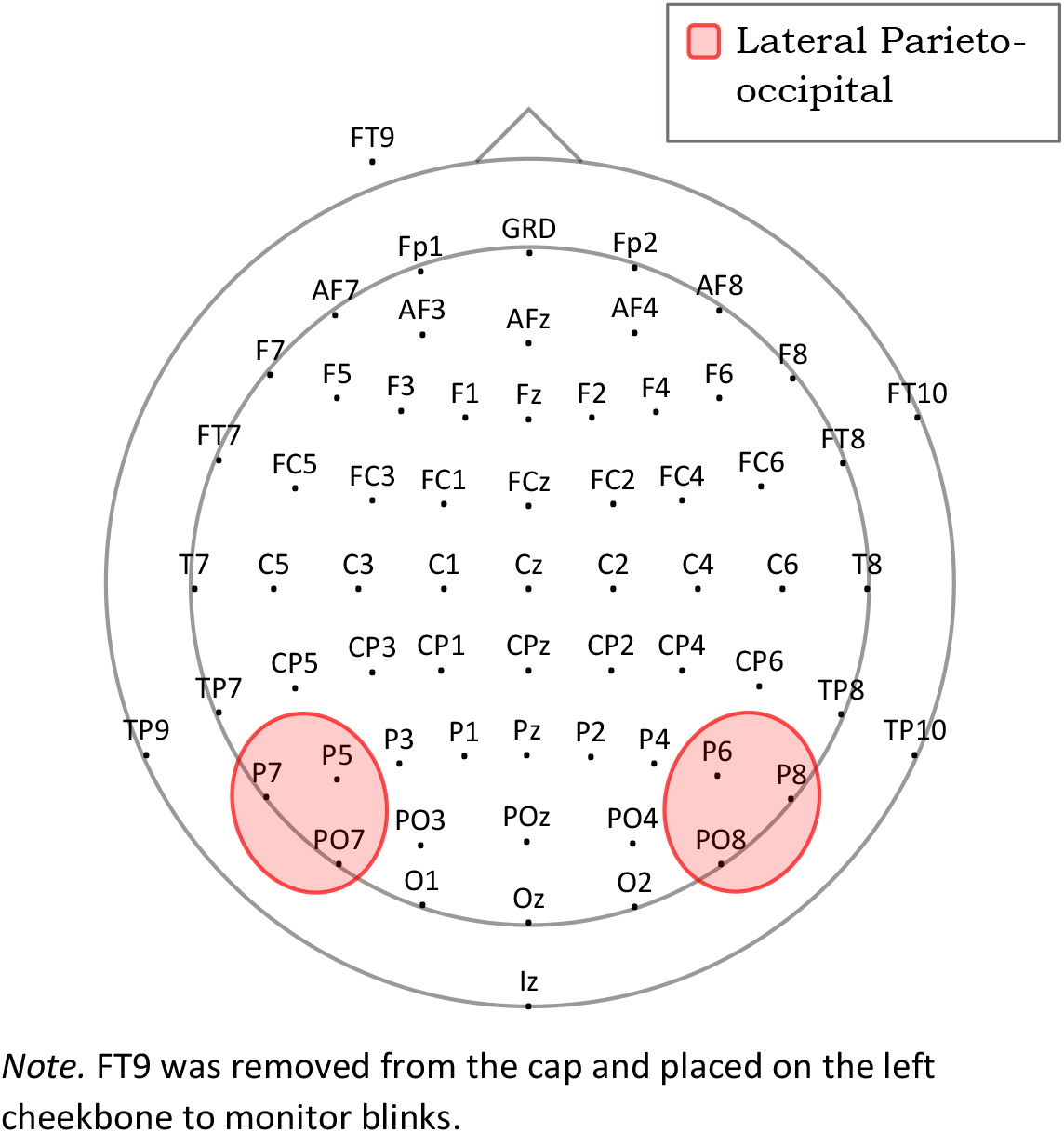
Map of Electrode PlacementIncluding the Lateral ROIs

Analysis was conducted on the mean amplitudes for the same time-period as before (175-250 ms) using a 2 (Hemisphere: Left; Right) x 2 (Congruency: Congruous; Incongruous) repeated-measures ANOVA. This revealed a main effect of Congruency, *F*(1, 23) = 5.81, *p* = .024, ƞp^2^ = .20, with more positive amplitudes for Incongruous (*M* = 4.78 µV) than Congruous (*M* = 4.26 µV) trials. The Hemisphere x Congruency interaction did not reach significance (*p* = .274). See Figure D2 for grand averaged ERPs.

**Figure D2.**
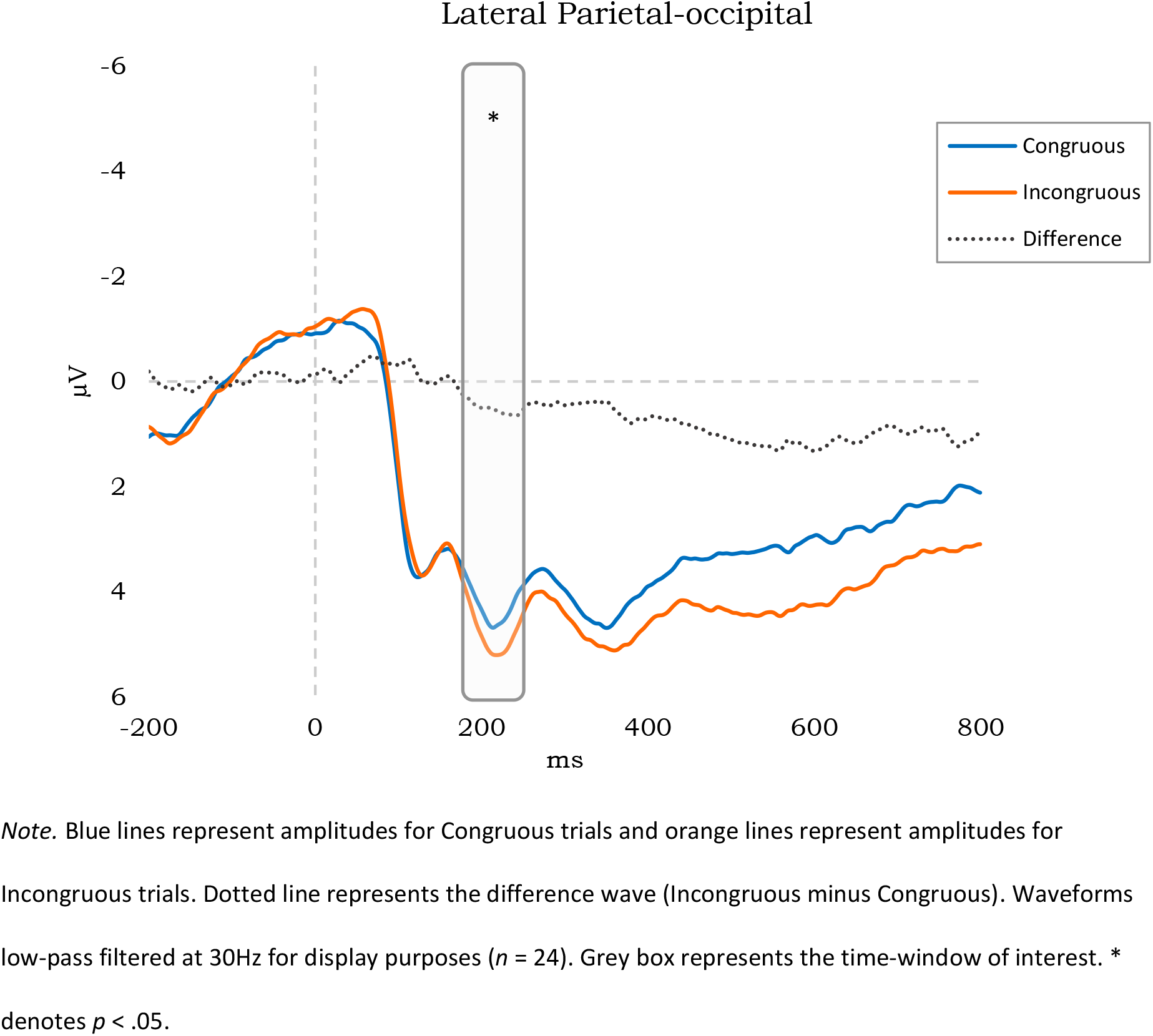
Grand-averaged ERPs for the Lateral Parieto-occipital Region, Collapsed Across Hemispheres

## Notes

We have no known conflict of interest to disclose

### Competing Interest Statement

The authors have declared no competing interest.

### Summary of Updates

A citation and associated clause has been removed from page 59.

